# One Scaffold – Different Organelles Sensors: pH-Activable Fluorescent Probes for Targeting Live Primary Microglial Cell Organelles

**DOI:** 10.1101/2021.05.05.442691

**Authors:** Krupal P. Jethava, Priya Prakash, Palak Manchanda, Harshit Arora, Gaurav Chopra

**Affiliations:** Department of Chemistry, Purdue University, West Lafayette, IN 47907, USA; Purdue Institute for Drug Discovery, West Lafayette, IN 47907, USA; Purdue Institute for Integrative Neuroscience, West Lafayette, IN 47907, USA; Purdue Institute for Inflammation, Immunology and Infectious Disease, West Lafayette, IN 47907, USA; Purdue Center for Cancer Research, West Lafayette, IN 47907, USA; Purdue University Integrative Data Science Initiative, West Lafayette, IN 47907, USA

**Keywords:** structure organelle relationship, pH-activable fluorescent probes, microglia, organelle targeting, lysosome

## Abstract

Targeting live cell organelles is important for imaging and to understand and control specific biochemical processes. Typically, fluorescent probes with distinct structural scaffolds have been used for targeting specific cell organelle. Herein, we aimed to design modular one-step synthetic strategies using a common reaction intermediate to develop new lysosomal, mitochondrial and nucleus targeting pH-activable fluorescent probes that are all based on a single boron dipyrromethane analogs. The divergent cell organelle targeting was achieved by synthesizing pH-activable fluorescent probes with specific functional groups changes to the main scaffold resulting in differential fluorescence and pKa. Specifically, we show that the functional group transformation of the same scaffold influences cellular localization and specificity of pH-activable fluorescent probes in live primary microglial cells with pKa’s ranging from ~4.5-6.0. We introduce a structure-organelle-relationship (SOR) framework targeting the nucleus (NucShine), lysosomes (LysoShine) and mitochondria (MitoShine) in primary mouse microglial cells. This work will result in future applications of SOR beyond imaging to target and control organelle-specific biochemical processes in disease-specific models.

---

Fluorescent organic materials have proven to be extremely useful for biological^1a^ and biomedical science.^1a^ Specifically, high-sensitivity fluorescent imaging of cellular organelles with enhanced spatial resolution allows direct visualization of dynamic cellular processes.^1b^ Small-molecule fluorescent probes are essential tools to monitor changes in biological processes in cellular organelles. These include imaging lysosomes,^2a-b^ mitochondria,^2c^ Golgi apparatus,^2d^ nucleus^2f^ among many others, and fluorescent probes are useful to track their abundance, localization, and function in cells. Lysosomes mainly act as a cellular ‘recycling plant’ to maintain intracellular and extracellular homeostasis^2a^ via the breakdown of carbohydrates, lipids, proteins, nucleic acids, cellular debris and other foreign pathogens. Mitochondria is the cellular ‘power plant’ that contains enzymes responsible for energy production needed for biochemical reactions and for energy metabolism to maintain cellular health. In addition, lysosomal and mitochondrial crosstalk is critical for cells and its dysfunction leads to diseases including neurodegeneration.^2e^ Often, imaging of cell organelles, irrespective of pH-activable property, involves specific fluorescent probes having different scaffolds that are prepared separately by multistep synthesis. In that context, a conceptual divergent synthetic strategy delivering a distinct organelle targeting from the same basic scaffold has remained elusive. Notably, if we could achieve the structure-organelle-relationship (SOR) of the same basic fluorescent scaffold, it would be beneficial in understanding the disease specific biological processes.

Microglia, the immune cells in the brain and macrophages in the periphery phagocytose (or engulf) extracellular material such as bacteria, virus, misfolded proteins, cell debris, etc. from their microenvironment into lysosomes for degradation.^3a^ Lysosomes are membrane-bound acidic organelles containing several enzymes (hydrolases, proteases, lipases, etc.) to actively breakdown the phagocytosed material.^3b^ Microglial cells are an excellent model for examining phagocytosis as well as lysosomal and mitochondrial activity *ex vivo* and *in vivo*^3c^ Microglia are the professional phagocytes in the brain that play a critical role in brain health and development.^3d^ It is known that microglial lysosomes are unable to effectively degrade a large quantity of phagocytosed amyloid-beta aggregates, which may be contributing to neurodegeneration during later stages of Alzheimer’s disease.^3e^ Furthermore, during neurodegeneration microglia releases dysfunctional mitochondria into their environment thereby exacerbating neuroinflammation.^3f^ It is, therefore, crucial to develop pH-activable chemical probes that target lysosomes and mitochondria to understand such cellular processes.^3c^ Fortunately, we can exploit the lysosomal acidic environment of pH 4.5 – 5.5^2a,4a^ to develop pH-activable fluorescent probes to visualize, track, and investigate lysosomal processes in live and fixed cells and *in vivo*.^3c^ For targeting mitochondria, we can exploit the negatively charged inner membrane of mitochondria to design a fluorescent probe with a positively charged functional group.^4b^ Importantly, the maintenance of a particular alkaline matrix (pH ~8) by pumping out protons dictates the normal physiological function of mitochondria.^4c^ However, during disease pathogenesis, impaired mitochondria undergo mitophagic elimination through lysosomal fusion.^2e^ Moreover, understanding the crosstalk between mitochondria and lysosomes using targeted fluorescent probes is important for the investigation of cellular processes leading to disease pathogenesis.^4d^ Nonetheless, if mitochondrial targeting fluorescent probe that has acidic pH-activable property, then such fluorescent probes could be useful to track mitochondrial fusion with acidic lysosomes.

The rational design of pH-activable florescent probes should satisfy several parameters: (i) ability to emit high fluorescence at lysosomal acidic pH and negligible fluorescence at cytosolic neutral pH, (ii) cellular permeability and uptake, (iii) non-specific binding to other cellular components, and (iv) good solubility. Several pH-activable fluorescent probes contain rhodamine,^5a^ coumarin,^5b^ napthalimide,^4b^ cyanine^4e-f^ and 4,4-difluoro boron dipyrromethane (known as BODIPY) based scaffolds.^5a^ The widely used are BODIPY-based scaffolds^5d^ due to a variety of synthetic routes to introduce diverse functionalities^6a-b^ for desired photophysical and spectroscopic properties. However, this process is not robust and minor changes in the substituents can significantly affect spectroscopic properties. Furthermore, if we can develop a synthetic strategy that could furnish divergent cell organelle targeting fluorescent probes showing SOR, it would not only reduce the chemical burden but also afford a convenient synthesis of fluorescent probes from a single synthetic intermediate.

Representative fluorescent probes with different chemical scaffolds that specifically target nucleus (Hoescht 33258),^2g^ lysosome (PhagoGreen)^6b^ or mitochondria (MitoTracker Green^TM^)^2f^ are shown in **Figure 1**. Here, we report a new modular design strategy for developing ratiometric BODIPY-based fluorescent probes targeting lysosomes, mitochondria and the nucleus that are highly fluorescent at acidic pH levels compared to cytosolic pH levels (**Figure 1**). During the course of the present study, we identified an interesting synthetic intermediate that is an excellent nucleus targeting fluorescent probe. We identified organelle targeting pH-activable fluorescent probes that are cell-permeable, and non-toxic to the cells. One of the most common starting materials to prepare BODIPY probes is 2,4-dimethyl-1*H*-pyrrole (**Scheme S1**, compound **1**) that exists as a liquid at room temperature. We used ethyl 2,4-dimethyl-1*H*-pyrrole-3-carboxylate (**2**) that is solid at room temperature, easy to handle, well-tolerated under reaction conditions, and it is still underrepresented in the literature to prepare boron dipyrromethene scaffold (**3**).^6c^ The additional functional group on the pyrrole ring system can serve as a handle for late-stage functionalization. Substitutions of BODIPY have significant effects on the excitation/emission property of a fluorescent probe but have only been studied at the 1,3,5,7-positions in scaffold **3**, using compound **1** but not compound **2** (**Scheme S1**).

**Figure 1.**
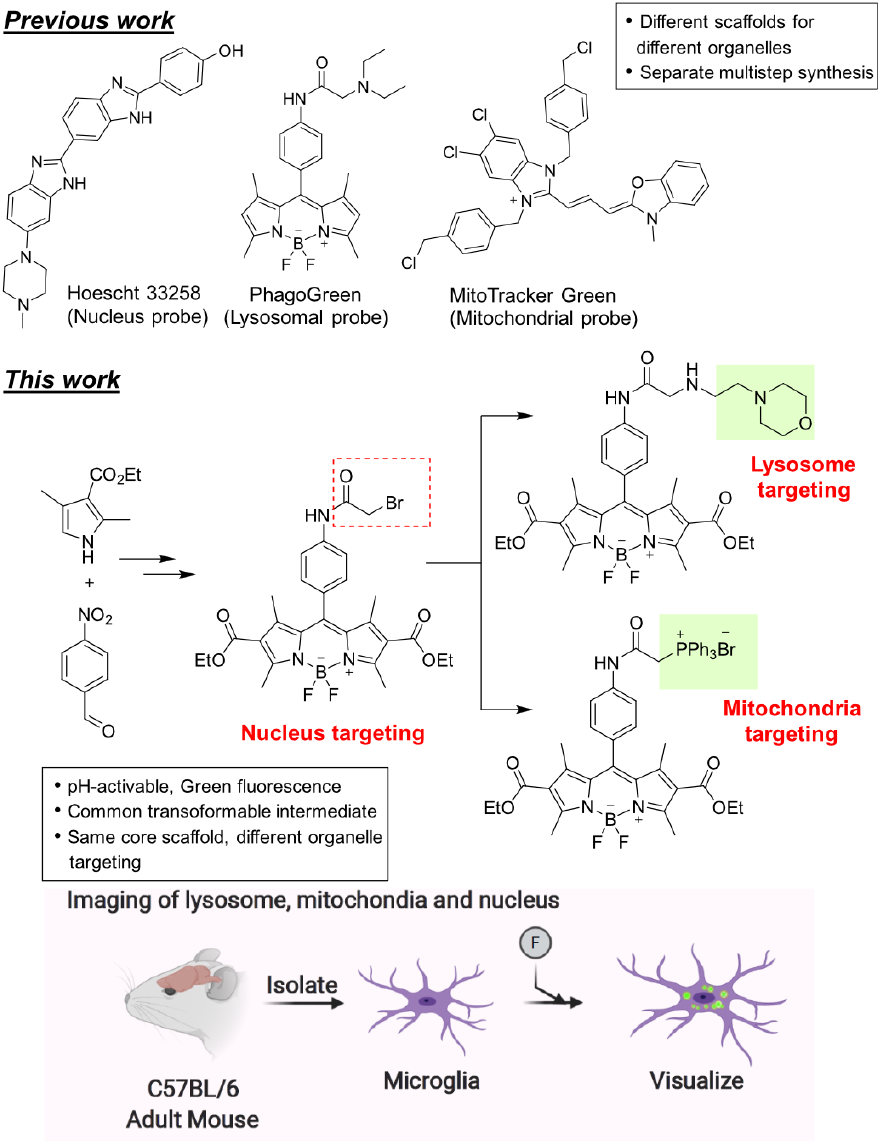
Our strategy to prepare cell organelle targeting probes in one synthetic scheme with a common scaffold.

**Scheme 1.**
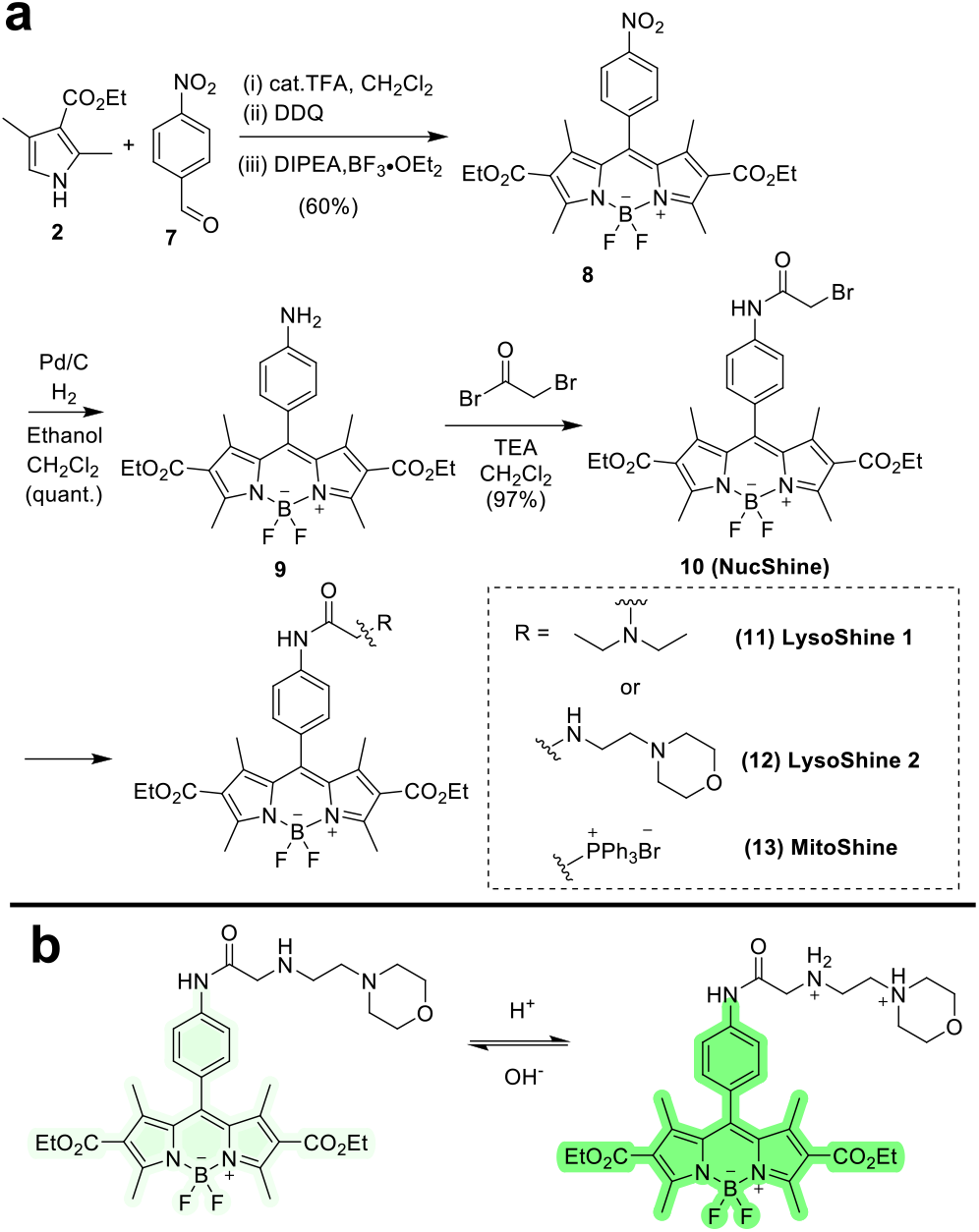
(**a**) Synthetic route for the synthesis of pH-activable NucShine (**10**), LysoShine probes (**11** and **12**) and MitoShine (**13**) probe; (**b**) A plausible response mechanism of the pH-activable probe (representative example)

Previous reports suggest that *N,N*-dimethylaniline functional group can be used to prepare pH-activable probes. Urano and Kobayashi et al.^7a^ and Kikuchi et al.^7b^ independently reported - NMe2 containing probe with pKa 4.3 and 4.5, respectively. Therefore, we strategically designed compound **5** to be synthesized in two steps (see **Scheme S2**) using 2,4-dimethyl-1*H*-pyrrole-3-carboxylate (**2**) and 4-(*N,N*-dimethylamine) benzaldehyde (**4**). Interestingly, the absorbance and fluorescence property of **5** showed the high fluorescence at pH 2.0 or less that was significantly different from similar probes reported in the literature (**Figure S1**).^7a-b^ This suggests that the ethylester functional group played a role in the pH-sensitive property of compound **5**. Notably, the Knoevenagel condensation between a methyl group at position 3 or 5 of BODIPY scaffold and substituted benzaldehyde was used to introduce extended conjugation to fine-tune the pH-sensitive property towards bathochromic (red) fluorescence shift. Specifically, compound **5** was reacted with 4-hydroxybenzaldehyde or 4-hydroxy 3-nitro benzaldehyde using piperidine, acetic acid as additives, and anhydrous toluene as a solvent under reflux condition to obtain expected conjugated products **6a-b** (**Scheme S2**).^7c,d^ However, **6a-b** were sparingly soluble in an aqueous medium and **6a** did not generate a fluorescence spectrum as solubility was affected by pH buffers. Compound **6b** was soluble in a mixture of DMSO: acetonitrile and showed pH-sensitivity with significant fluorescence at pH less than 6 (**Figure S2**) due to the presence of a nitro group at *ortho*-position to the hydroxyl group.^7e^ However, poor solubility of these compounds does not warrant its use in primary cells and for future *in vivo* applications.

To overcome these challenges, we designed another synthetic route to achieve a facile and modular synthesis of pH-activable fluorescent probes targeting lysosomes, mitochondria and the nucleus (**Scheme 1a**). Herein, we envisioned preparing an important synthetic intermediate that can be transformed into different organelle targeting probes in one step. Using the common synthetic intermediate, we installed functional groups that target a specific cell organelle individually. For example, we planned to synthesize pH-responsive lysosome targeting probes, using morpholine as a lysosome-targeting moiety. ^1a,5c,8a-b^ The BODIPY fluorophore can be fine-tuned by the photo-induced electron transfer (PET) mechanism of the lone pair electrons of a nitrogen atom in the morpholine as well as secondary amine functional group (**Scheme 1b**). A similar mechanism can be envisioned when the diethylamine group is present. We started the synthesis of designed pH-responsive probes by the reaction of compound **2** with 4-nitro benzaldehyde (compound **7**) afforded compound **8** as a brownish-black solid. The nitro compound **8** was reduced successfully into amine-containing compound **9** using Pd/C in ethanol: CH_2_Cl_2_ solvent mixture. Next, compound **9** readily reacted with bromo acetyl bromide to give **10** with excellent yields. The – CH_2_Br handle provides easy access to substitute with an amine functional group-containing reactant to get the final product. We used diethylamine and 2-aminoethyl morpholine substrate to synthesize compounds **11** (**LysoShine 1**) and **12** (**LysoShine 2**), respectively. We noticed that – CH_2_Br in intermediate **10** could serve as a useful synthon to introduce another targeting moiety such as cationic triphenylphosphine for mitochondria targeting. Therefore, we prepared a mitochondrial probe (compound **13**) also using intermediate **10** when reacted with triphenylphosphine under the inert condition and named compound **13** as **MitoShine**.

Next, we investigated the absorption and fluorescent properties of LysoShine 1, LysoShine 2, and MitoShine (**Figure 2**). The LysoShine 1 in different pH solutions of phosphate buffer (1% DMSO in 1M PBS) has an absorption centered around 500 nm and emission maximum at 505 nm with 480 nm excitation. LysoShine 1 is highly fluorescent at pH 4 compared to reduced fluorescence that gradually decreased from pH 5 to 7 (pKa of 5.4, **Figure 2**). On the other hand, LysoShine 2 (max. absorption 500 nm, max. emission 512 nm, excitation 480 nm) showed better pH-sensitivity with a gradual increase for pH less than 6 (pKa of 6.0). We tested the mitochondrial targeting property of MitoShine (max. absorption 505 nm, max. emission 535 nm, excitation 480 nm). Interestingly, MitoShine showed significantly higher fluorescence at pH 4 compared to other pH values (**Figure 2**) with a pKa of 4.4 suggesting possible use to image mitochondria-lysosome crosstalk. We also tested the photophysical property of the important synthetic intermediate compound **10** (absorption maximum 505 nm, max, emission 510 nm at excitation 480 nm, pKa 3.2, **Figure S3**). Unlike, compound **6a-b**, LysoShine 1 and 2, MitoShine maintains its solubility upon addition of pH solutions (final DMSO concentration 1%), so we considered it as compatible for further biological experiments with cells.

**Figure 2.**
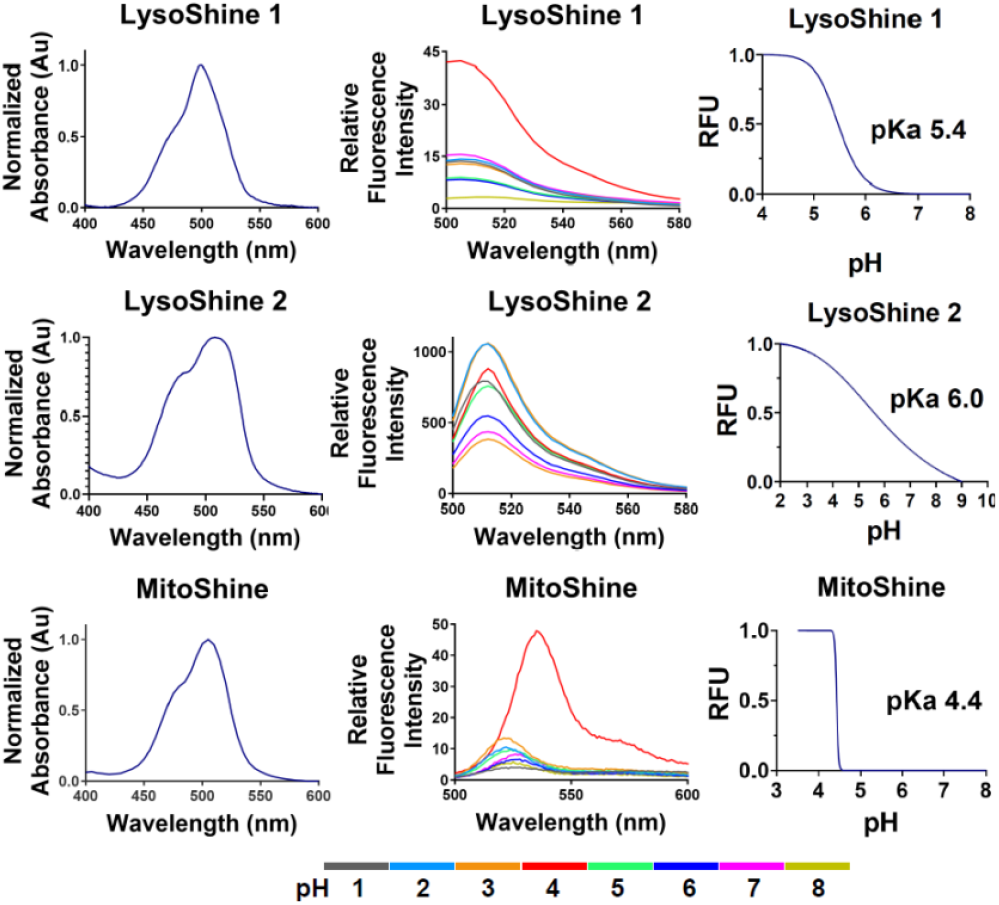
Absorption, Fluorescence spectrum of compound **LysoShine 1**, **LysoShine 2**, **MitoShine** at different pH solutions and their pKa values. Spectrum recorded at room temperature in PBS buffer at varying pH with 1% (v/v) of DMSO, in all cases probe concentration = 10 μM.

Next, we performed a series of experiments with BV2 microglia cell line and with primary microglia isolated from mouse brains towards applications in biological systems. We performed MTT assay with 1, 5 and 10 μM concentrations of the probes at 24 hours with BV2 microglia to measure the effect on metabolic activity (**Figure S4**). All the compounds maintained 80% or higher cellular metabolic activity except for MitoShine, which maintained a high cellular metabolic activity at 1 μM but only 10% activity at 5 μM or higher concentrations. We also assessed the cytotoxicity of the probes on BV2 using the lactate dehydrogenase (LDH) assay that measures the levels of LDH released by dying cells into the cell culture medium. In 2 hours assay, LysoShine 1 and 2 did not show any cytotoxicity at any tested doses. Contrarily MitoShine showed no cytotoxicity at 1 and 5 μM, it showed less than 10% cytotoxicity at 10 μM (**Figure S5a**). In 24 hours treatment, LysoShine 1 and 2 showed less than 10% cytotoxicity at all the tested doses compared to MitoShine that showed around 20% cytotoxicity at higher doses of 10 μM and 5 μM (**Figure S5b**). Therefore, the lower metabolic activity of the cells at higher concentrations of MitoShine for 24 hours may lead to cytotoxicity. For a fluorescent probe, it is essential to have high cell permeability. So, we also checked the uptake efficiency of synthesized probes. In 2 hour treatment, almost all probes showed uptake efficiency around 80% or higher at all doses except for MitoShine that showed 30% of uptake at 10 μM (**Figure S6**). Next, having important information (fluorescent property, cytotoxicity and uptake efficiency) in hand, we focused on the cell imaging experiments and flow cytometry analysis to test these pH-activable probes in primary microglia. Notably, compound **10** is one of the important intermediate and found to be cell permeable (**Figure S6**) as well as not cytotoxic (**Figure S4**) to BV2 cells. Interestingly, MitoTracker Green has a -CH_2_Cl functional group while compound **10** has -CH_2_Br. Whereas previously reported microglia specific probes have -CH_2_Cl group.^7f^ So, it would be interesting to test the behavior of compound **10** inside the cell. So, first, we checked the localization of compound **10**, which showed high fluorescence at pH less than 3 (**Figure S3)**. Interestingly, in the confocal imaging, we observed bright green puncta that colocalized with nuclei staining dye DAPI (**Figure S8**). For nucleus targeting, DAPI and Hoechst dyes are the most widely used dyes in the field and only a few novel nucleus targeting probes have been reported due to the challenges of nucleus targeting such as poor target efficiency, membrane impermeability etc.^2f-g^ The compound **10** (named **NucShine**) satisfy these requirements and potentially a new nucleus targeting probe. These observations motivated us to explore–how transforming one functional group to another impacts the cell organelle targeting ability of these pH-activable probes in primary mouse microglia.

We performed flow cytometry analysis to evaluate the intensity of the fluorescent signals of the probes in primary mouse microglia (**Figure 3a**) for future cell sorting applications. The cells treated with the fluorescent probes at 10 μM for 2 hours showed increased green fluorescence compared to the cells treated with the vehicle only (unstained control) thereby clearly discriminating between the probe-treated and untreated cells (**Figure 3b-d**). Furthermore, LysoShine 1 showed higher fluorescent intensity than LysoShine 2 within the cells (**Figure 3** along with the commercially available LysoTracker dye, or with (ii) MitoShine probe and the MitoLite dye, we were able to identify over 95% of the live cells that are LysoShine^+^LysoTracker^+^ and around 87% of live cell subset that was MitoShine^+^MitoLite^+^ (**Figure S7**). The ability to identify the probe-specific individual cells also demonstrates the possibility of sorting the cell subsets for downstream analysis in the future.

**Figure 3.**
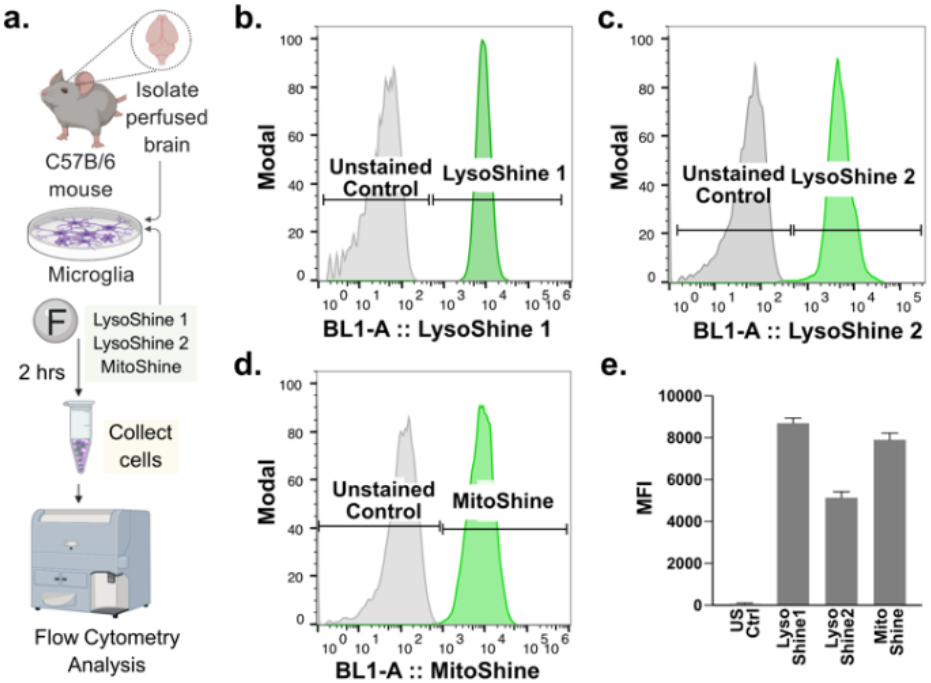
**(a)** Schematic for flow cytometry analysis in live cells. **(b-d)** Demonstrates the fluorescence of LysoShine 1, LysoShine 2, and MitoShine upon uptake by primary mouse microglia (live cells). Modal corresponds to a percentage of the maximum count. **(e)** Median fluorescence intensity (MFI) values for each probe. US Ctrl is unstained control. Gating strategy and flow plots with commercial dyes in supporting information.

Next, to confirm the localization of the fluorescent probes within the lysosomal organelles, we performed confocal imaging of the probes with primary microglia (**Figure 4a**). The green fluorescent signal of LysoShine 1 and LysoShine 2 appeared as several bright puncta around the nuclei as well as in the cytosolic regions of the cells. The localization of the probes into the intracellular acidic organelles was confirmed by co-staining with the commercially available LysoTracker Red DND-99 dye (**Figure 4b-c**). We obtained similar fluorescence in BV2 microglia (**Figure S9**). The LysoShine probes clearly co-localized with the LysoTracker dye within the acidic lysosomes and not with DAPI-labelled nuclei, confirming targeting in cells.

**Figure 4.**
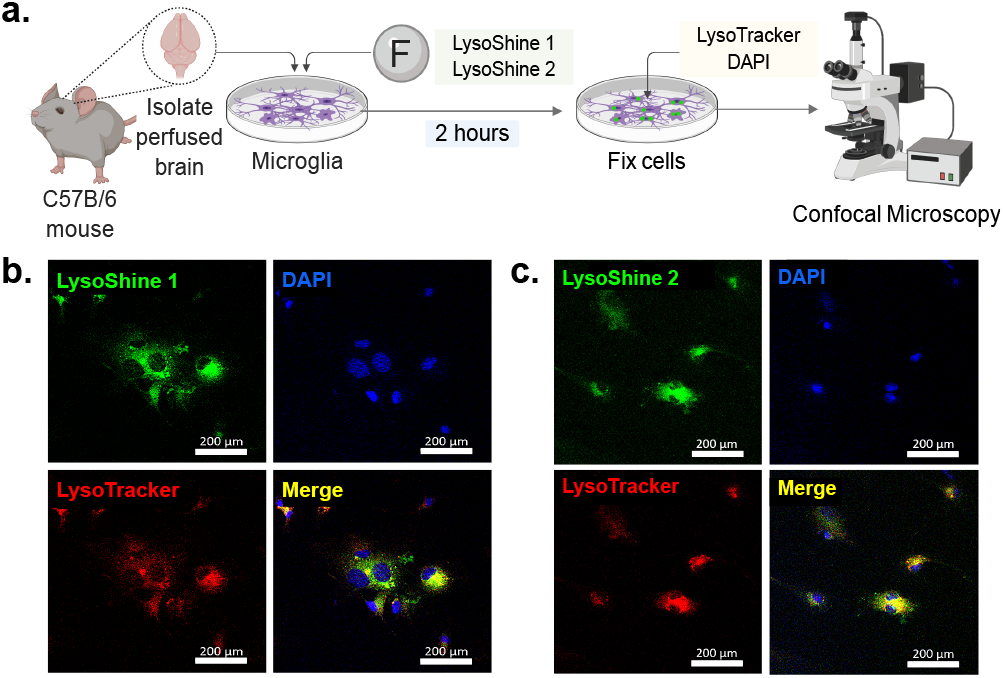
**(a)** Experimental design for fluorescence imaging of primary mouse microglial cells with the **(b)** LysoShine 1 and **(c)** LysoShine 2 (green). The localization of the compounds in the lysosomal acidic compartments is shown with the LysoTracker dye (red). Nuclear DNA is stained with DAPI (blue). Scale bars represent 200 μm.

The localization of the mitochondrial probe, MitoShine, was similarly evaluated in primary microglia with confocal imaging. The cells showed a bright green fluorescent signal and the green puncta with MitoShine within the cells clearly co-localized with MitoLite, a commercial mitochondrial dye (**Figure S10a**). Interestingly, we observed several other regions (green puncta) labeled with MitoShine that did not overlap with MitoLite (**Figure S10b**) suggesting that localization of the MitoShine probe may be localizing in other organelle along with the mitochondria. Interestingly, in a separate experiment, we observed an overlap of the MitoShine probe with LysoTracker, an acidic lysosomal dye (**Figure S10c**).

We asked if MitoShine could be used to study mitochondrial transport and recycling as a means to dynamically monitor organelle quality^4d^ or identify mitophagic elimination through lysosomal fusion.^2e^ We therefore treated the primary microglia with MitoShine for 2 hours and later with both LysoTracker and MitoLite at the same time. Confocal imaging demonstrated the colocalization of MitoShine in both mitochondrial as well as in lysosomal organelles (**Figure 5**). In addition, overlapping both channels of LysoTracker and MitoLite dyes indicates the merging of mitochondria and lysosomes in these cells (**Figure S11**) suggesting the possible use of MitoShine for mitochondrial-lysosomal fusion processes for future biological applications.

**Figure 5.**
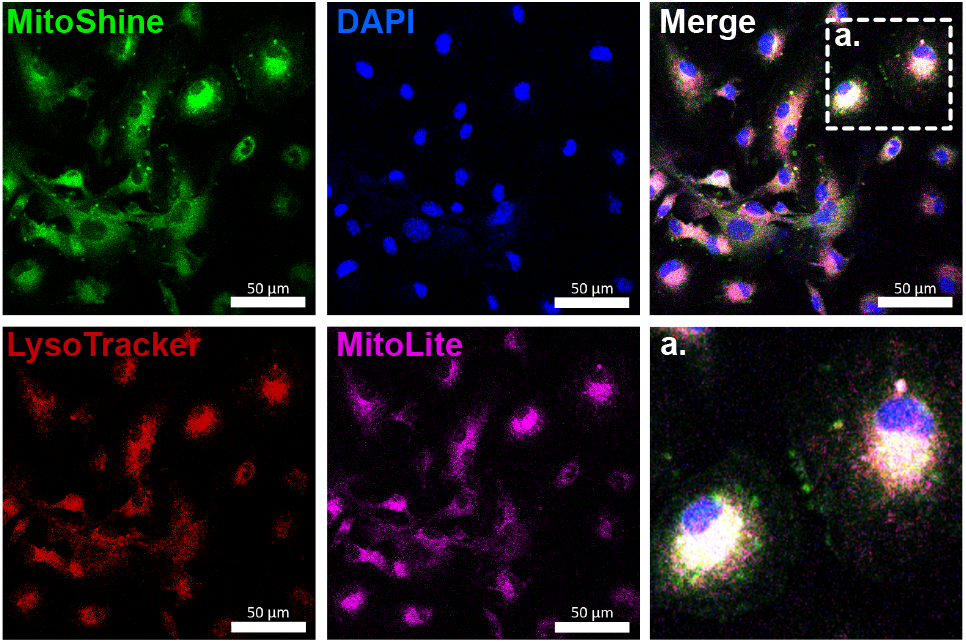
Fluorescence imaging of primary mouse microglial cells with MitoShine. Localization of the compound was observed in mitochondria (magenta, MitoLite dye) as well as in the acidic lysosomal organelles (red, LysoTracker dye). **(a)** Magnified image with nuclear DNA stained with DAPI (blue). Scale bar 50 μm.

In summary, we have developed a modular synthetic strategy for pH-activable fluorescent BODIPY probes to achieve divergent targeting of cellular organelles that are tested in primary cells and available for future biological use *in vivo*. We showed how the transformation of a critical synthetic intermediate containing bromomethyl group into various derivatives results in a distinct targeting ability of the probe affording SOR for lysosomal, mitochondrial and nucleus targeting probes. The synthesized fluorescent probes have high fluorescence at acidic lysosomal pH compared to cytosolic neutral pH. The pH-activable property of fluorescent probes was utilized for targeting lysosomes and mitochondria in primary mouse microglial cells. Besides, the synthetic intermediate or the probes with free amine group can also be used for several bioconjugation reactions, including targeting a protein of interest such as the Aβ(1-42) peptide^3c,8c^ to investigate target-specific microglial uptake towards specific cellular organelles. Further derivatization of pH-activable probes and biological applications will be explored in the future.

## ASSOCIATED CONTENT

Copies of ^1^H and ^13^C NMR spectra for all new compounds, supporting text for the methods and protocols, supporting figures for biological experiments.

## AUTHOR INFORMATION

### Author Contributions

All authors have approved the final version of the manuscript.

### Notes

Any additional relevant notes should be placed here.

## ACKNOWLEDGMENT

This work was supported, in part, by Purdue Research Foundation award, Purdue Integrative Data Science Institute award, NIH NCATS ASPIRE Design Challenge awards and the United States Department of Defense USAMRAA award # W81XWH2010665 to Gaurav Chopra. Additional support, in part by, the Stark Neurosciences Research Institute, the Indiana Alzheimer Disease Center, Eli Lilly and Company, the Indiana Clinical and Translational Sciences Institute grant # UL1TR002529 from NCATS, and the Purdue University Center for Cancer Research funded by NIH grant # P30 CA023168 are also acknowledged.

## ABBREVIATIONS

BODIPY: 4,4-difluoro boron dipyrromethane
DMSO: dimethyl sulfoxide
MTT: Microculture Tetrazolium
LDH: Lactate dehydrogenase

---

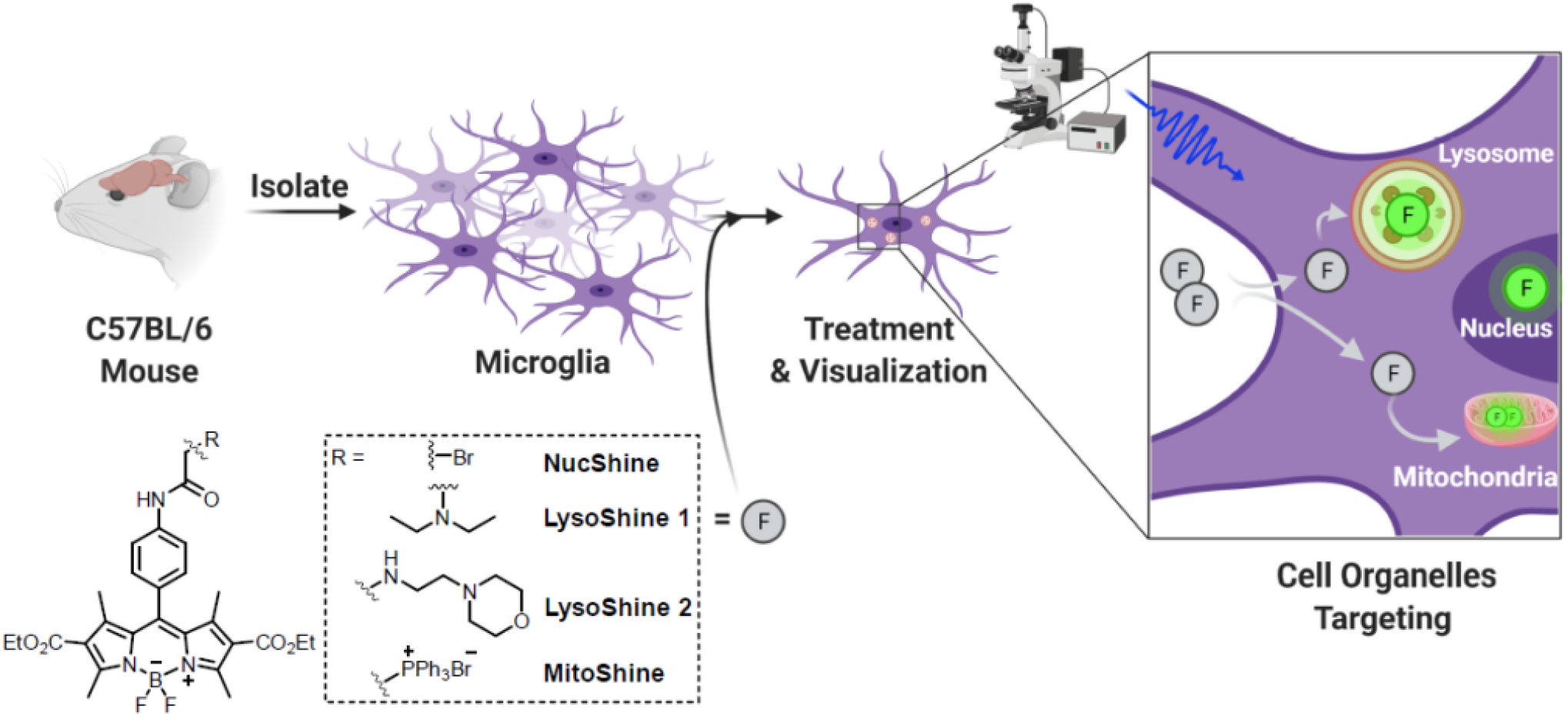

