## Supporting Information for "One Scaffold – Different Organelles Sensors: pH-Activable Fluorescent Probes for Targeting Live Primary Microglial Cell Organelles"

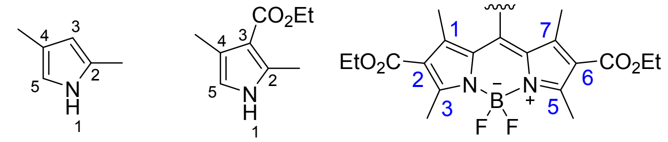

2,4-Dimethyl-1*H*-pyrrole (**1**)

Ethyl 2,4-dimethyl-1*H*-pyrrole-3-carboxylate (**2**)

2,6-Dicarboxylate substituted BODIPY (**3**)

**Scheme S1**. Strategy to synthesize BODIPY scaffold 3 with ethylester functional group

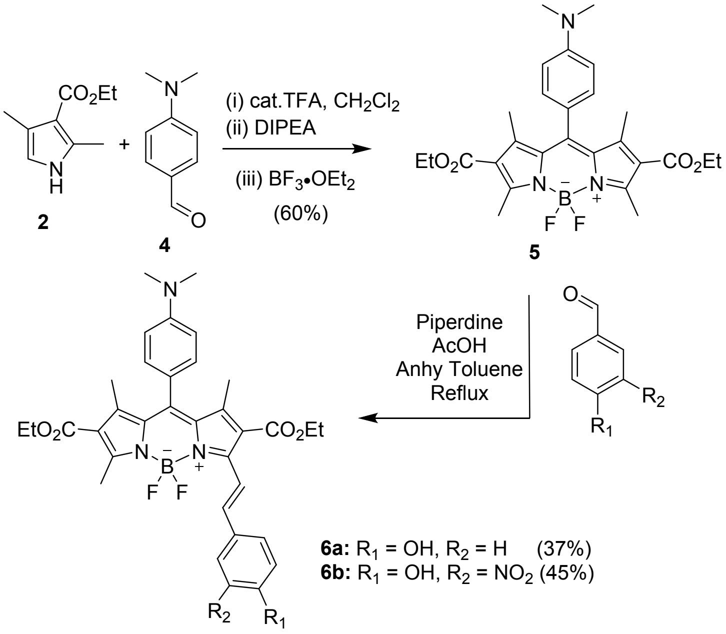

**Scheme S2**. Synthetic route for the approach towards the synthesis of pH-activable probes with extended conjugation

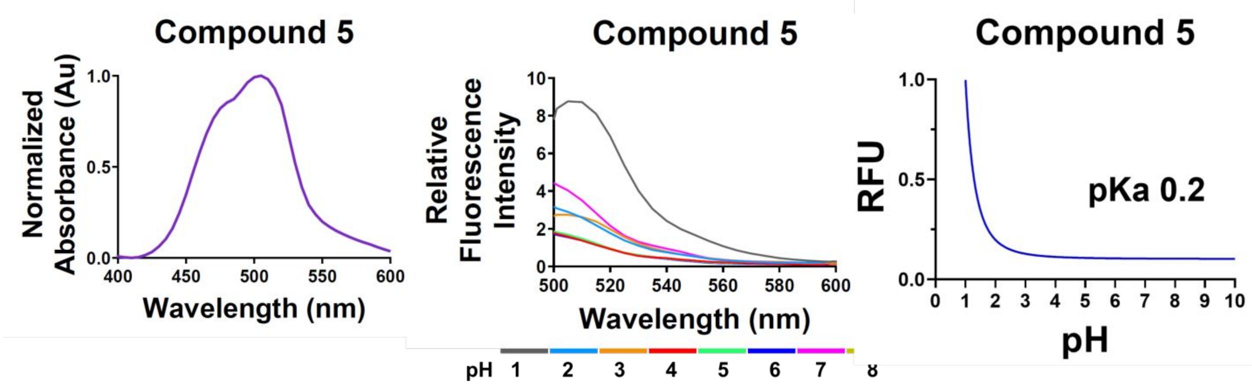

**Figure S1.** Absorption and fluorescence spectrum of compound **5** at 10 µM concentration in PBS (1% DMSO). The absorption maximum 505 nm, excitation/emission 480/512 nm, pKa 0.2 were observed. The pH solutions were prepared in 1M PBS buffer using dilute sodium hydroxide or hydrochloric acid.

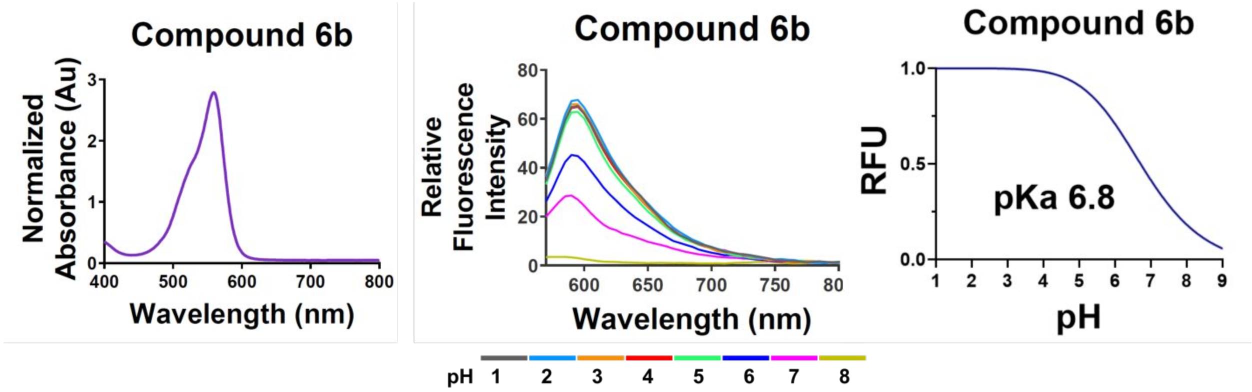

**Figure S2.** Absorption and fluorescence spectrum of compounds **6b** at 10 µM concentration in PBS [20% DMSO:MeCN (1:1 mixture)]. The absorption maximum 560 nm, excitation/emission 550/590 nm, pKa 6.8 were observed. The compound **6a-b** were sparingly soluble in the aqueous medium, so the absorbance and fluorescence spectrum were recorded using a mixture of phosphate buffer containing 20% DMSO:MeCN (1:1). No florescent spectrum was recorded for compound **6a** as it was not soluble and precipitated out after addition of pH solutions. pH solutions were prepared in 1M PBS buffer using dilute sodium hydroxide or hydrochloric acid.

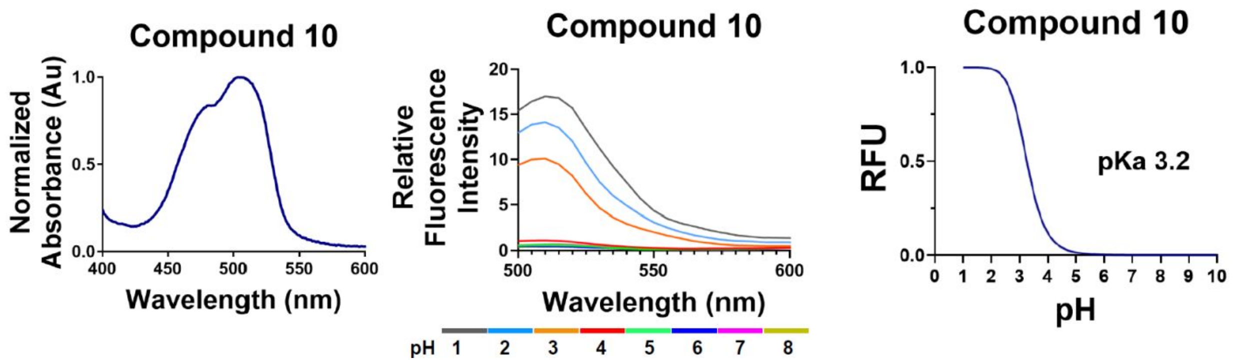

**Figure S3.** Absorption and fluorescence spectrum of compound **10 (NucShine)** at 50 μM concentration in PBS (2% DMSO). The absorption maximum 505 nm, excitation/emission 480/510 nm, and pKa 3.2 were observed. The pH solutions were prepared in 1M PBS buffer using dilute sodium hydroxide or hydrochloric acid.

### BIOLOGICAL EXPERIMENTAL SECTION

#### Animals

All mice were handled according to the Purdue Animal Care and Use Committee (PACUC) guidelines. Adult C57BL/6 mice (5-7 months old) bred in house were used for isolating microglia.

#### Culture and maintenance of BV2 microglia

BV2 mouse microglia cells were generously gifted by Dr. Linda J. Van Eldik (University of Kentucky, USA). The BV-2 cell line was developed in the lab of Dr. Elisabetta Blasi at the University of Perugia, Italy. Cells were maintained at 37 °C and 5% $CO_2$ in DMEM/Hams F-12 50/50 Mix supplemented with 10% Fetal Bovine Serum (FBS), 1% L-Glutamine, and 1% Penicillin/Streptomycin. For imaging experiments, 70,000 cells/2 mL/well were seeded on glass coverslips (Corning #12-553-450) in 6-well plates (Corning #08-772-1B). For flow cytometry experiments, 20,000 cells/0.5 mL were seeded in 24-well plates (Corning #3526).

#### Primary Mouse Microglia Isolation and Culture

A detailed protocol for the isolation and culture of primary microglia from adult mouse brains is previously described[2]. Briefly, $CD11b^+$ primary microglia were isolated from adult C57BL/6 mice aged 5-7 months of age (both male and female sexes) and cultured as follows. Mice were euthanized with $CO_2$ following the PACUC guidelines, perfused brains were removed and cut into small pieces before homogenizing them in DPBS++ with 0.4% DNase-I on the tissue dissociator at 37 °C. After filtering the cells through a 70 μm filter, myelin was removed two times, first using gradient centrifugation with Percoll PLUS reagent followed by the use of myelin removal beads on the magnetic column separators. After myelin removal, $CD11b^+$ cells were selected from the single cell suspension using the $CD11b^+$ beads as per the manufacturer's instructions. The $CD11b^+$ cells were finally resuspended in microglia growth media, further diluted in TIC (TGF-β, IL-34, and cholesterol containing) media containing 2% FBS before seeding $0.1x10^6$ cells/500 μL/well of a 24-well plate. The cells were maintained at 37 °C and 10% $CO_2$ with half-media change every other day until the day of compound treatment (around 12-14 div). For confocal imaging experiments, around 50,000 cells/2mL cells were sub-cultured at the center of 35mm glass-bottom imaging dishes (FluoroDish™ #FD35).

#### Reconstitution of fluorescent probes in DMSO and cell treatment

The dried fluorescent probes powders were resuspended in cell grade DMSO to prepare 5 mM stock solutions. This stock was used to make a 1, 5, or 10 μM dilution of the probe in cell culture media. The diluted probe solutions were filtered through 0.22 μm filters before adding to the cells.

#### Determination of Metabolic Activity of BV2 microglia with MTT assay

Murine microglial BV2 cells (10,000 cells/well) were seeded in a 96 well plate and cultured for 24 hours at 37 °C in a 5% $CO_2$ incubator. Next day, the media was aspirated, and the cells were rinsed twice with PBS (pH 7.4) followed by treatment with 1, 5, or 10 μM of the probe solution made in DMSO (final DMSO concentration = 0.05%) for 24 hours in the incubator. Next, the media was aspirated and 20 μL of 5 mg/mL 3-(4,5dimethylthiazol-2-yl)-2,5-diphenyltetrazolium bromide (MTT) solution was added to the cells and then incubated for three hours. The MTT solution was then removed and 100 μL DMSO was added to dissolve the violet formazan crystals. The plate was shaken for 5 minutes using an orbital shaker. The absorbance value at 450 nm was recorded and the percent cell metabolic activity was calculated as the ratio of sample absorbance to control absorbance.

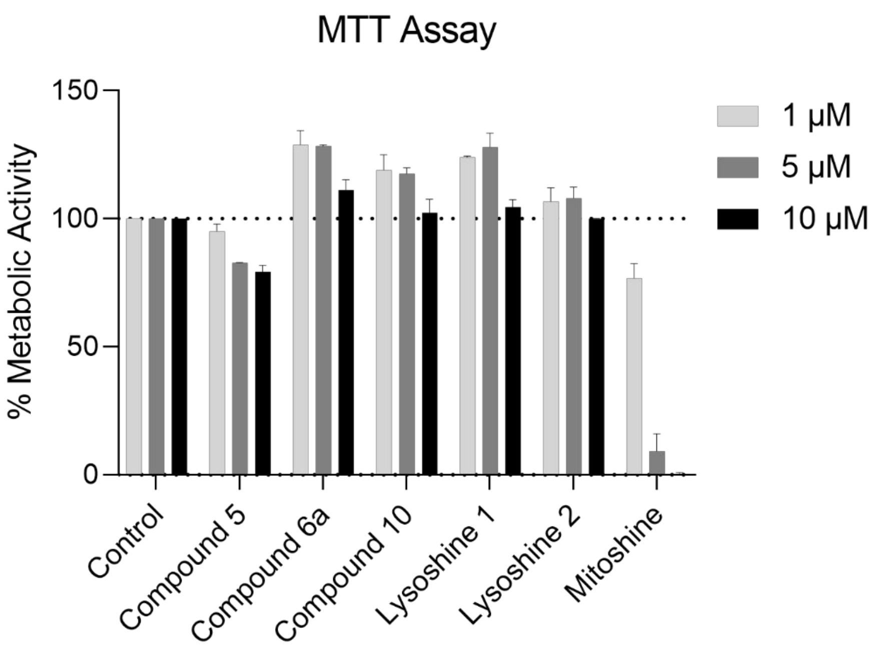

**Figure S4.** Effect of the fluorescent probes at 1, 5 or 10 μM concentration on the metabolic activity of BV2 microglia. BV2 microglia were treated with different concentration (1, 5, 10 μM) of the florescent probes for 24 hours prior to performing the MTT assay. Bars depict n=3 data with SD. Effect of the probes on the metabolic activity is calculated relative to the DMSO controls set to 100% metabolic activity.

#### Determination of cytotoxicity of the probes on BV2 microglia with LDH assay

The cytotoxicity of the fluorescent probes was determined using the Invitrogen™ CyQUANT™ LDH Cytotoxicity Assay kit which measures the release of Lactate dehydrogenase (LDH) from the dead and dying cells. The assay was performed per the manufacturer's protocol. Briefly, BV2 cells (5000 cells/100 μL/well) were seeded in a 96 well plate and cultured for 24 hours at 37 °C in a 5% $CO_2$ incubator. After 24 hours, the media was removed, and the cells were treated with 1, 5, or

10 μM of the probes for 2 or 24 hours of incubation. Three additional wells were treated with the given lysis buffer for 45 minutes (positive control). After the corresponding incubation period, the total LDH release was measured using a fluorescent plate reader. The cells without any probe treatment correspond to spontaneous LDH release and were taken as negative control and the cells treated with the lysis buffer correspond to maximum LDH release. The percentage cytotoxicity of the probes was determined as follows:

$$\% \text{ cytotoxicity} = \frac{[\text{Probe-treated LDH activity} - \text{Spontaneous LDH activity}] \times 100}{[\text{Maximum LDH activity} - \text{Spontaneous LDH activity}]}$$

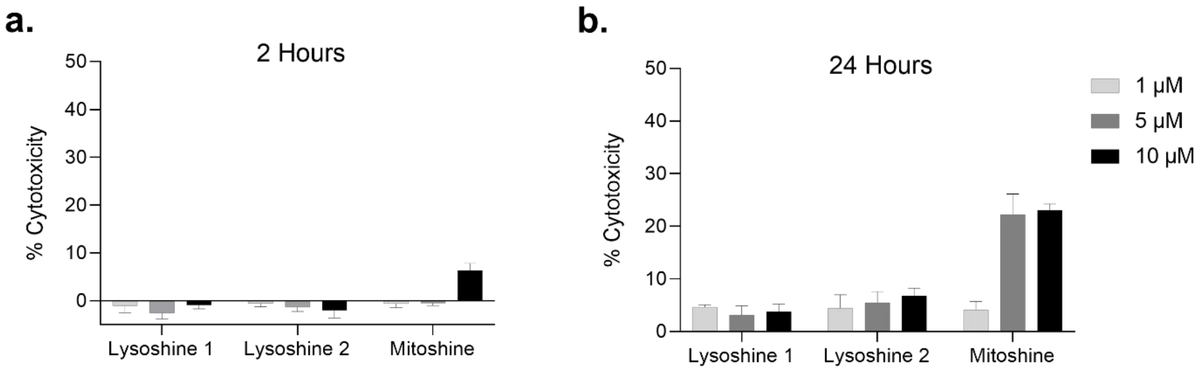

**Figure S5.** Percent cytotoxicity of the LysoShine 1, LysoShine 2, and MitoShine fluorescent probes. Percent (%) cytotoxicity after **a**) 2 hours treatment, **b**) 24 hours treatment with the probes was calculated relative to % cytotoxicity of maximum LDH positive control that was set to 100% (not shown) as described in the protocol above. Bars depict n=2 data with SD.

#### Determination of Cellular Uptake of the Fluorescent Probes

Cellular uptake of the florescent probes was determined *in-vitro* as described previously.[3] Briefly, BV2 cells (20,000 cells/well) were seeded in a 96 well plate and cultured overnight at 37 °C in a 5% $CO_2$ incubator. Next day, the media was removed, and the cells were treated with 1, 5, or 10 μM of the probes for 2 hours. After incubation, the supernatant was collected, and the absorbance was recorded at the respective absorbance maxima of the probe. This was compared with absorbance of the probe in media without cells. The percent cellular uptake was calculated as percentage of the ratio of absorbance of the supernatant to the absorbance of probe solution without cells:

$$\% \text{ Uptake Efficiency} = 1- \frac{[A_{\text{supernatant}}] \times 100}{[A_{\text{probe}}]}$$

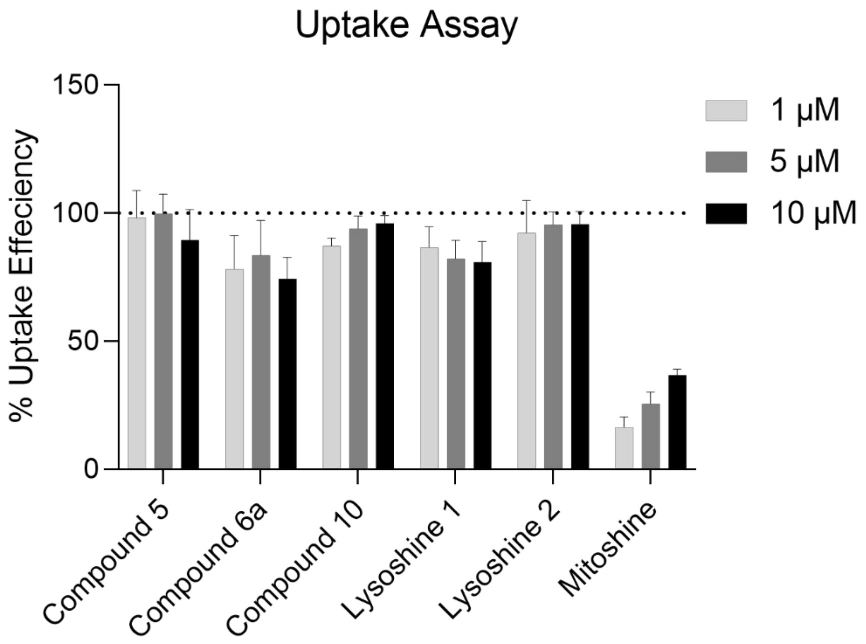

**Figure S6.** Percent uptake efficiency of the fluorescent probes in BV2 microglia. Cellular uptake of the probe at different concentrations after two hours of incubation. The % uptake efficiency was determined as the percentage of probe taken up by cells out of the total amount of probe in the initial incubation solution. Bars depict n=3 data with SD.

#### Flow cytometry analysis

Primary mouse microglia were treated with 1, 5, or 10 μM fluorescent probes for 2 hours for the cells to uptake the probe. The media was then aspirated, and the cells were incubated with 100 nM LysoTracker (1 mM stock from Thermo #L7528) or with 0.5x MitoLite (1000-fold dilution from 500x stock of AAT Bioquest #22678) for 1.5 hours. The media was aspirated and 500 μL/well of cold (4 °C) phosphate buffered saline was added to the cells. The cells were gently detached from the plates and transferred to corresponding vials. Finally, 0.1 ug/mL of DAPI was added to the suspended cells (500 μL volume) for 3 mins before analyzing the cells on the Attune NxT flow cytometer (Invitrogen). All the cells were first gated on the side and forward scatter plot (SSC vs. FSC) followed by gating the live cells using the DAPI nuclear stain. Around 10-20 thousand cells were gated from the live cells in order to analyze the cellular fluorescence signal corresponding to the LysoShine/Lysotracker or Mitoshine/MitoLite probes.

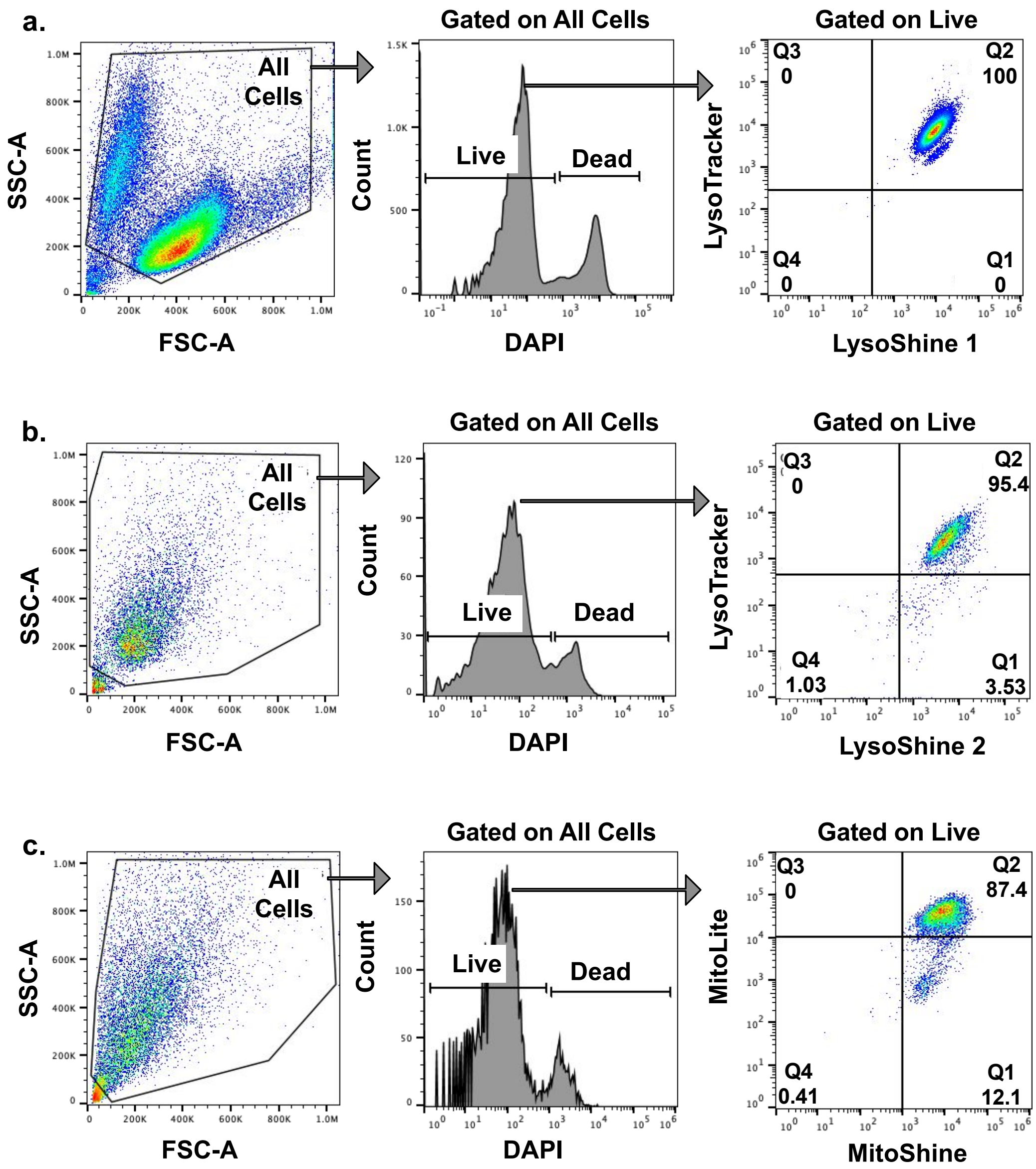

Figure S7. Gating strategy for flow cytometry analysis of primary mouse microglia treated with the fluorescent probes. All cells selected in the SSC-A vs FSC-A plots were used to visualize and quantify live and dead cells stained with DAPI dye. From this, live cells were selected to identify and quantify the (**a, b**) LysoTracker$^+$LysoShine$^+$ cells or (**c**) MitoLite$^+$MitoShine$^+$ cells.

#### Confocal imaging

The localization of the fluorescent probes were visualized using confocal microscopy. The cells were incubated with 10 μM probe for 2 hours and then the media was aspirated. The cells were then incubated with 100 nM LysoTracker (1 mM stock from Thermo #L7528) or with 0.5x MitoLite (1000-fold dilution from 500x stock of AAT Bioquest #22678) for 1.5 hours. The media was aspirated, and the cells were fixed with 4% paraformaldehyde for 10 mins followed by nuclear staining with 1 μM/mL DAPI for 5 mins. For BV2 microglia grown on glass coverslips, the coverslips were removed from the wells and transferred to glass slides with a drop of anti-fade reagent (Thermo Fisher Scientific #P36930). For primary mouse microglia grown in 35 mm glass-bottom dishes, the cells were treated with a few drops of the anti-fade reagent and taken for imaging. The images were captured on a Zeiss LSM 880 Upright Confocal microscope equipped with Plan-Apochromat 20x/0.8 objective.

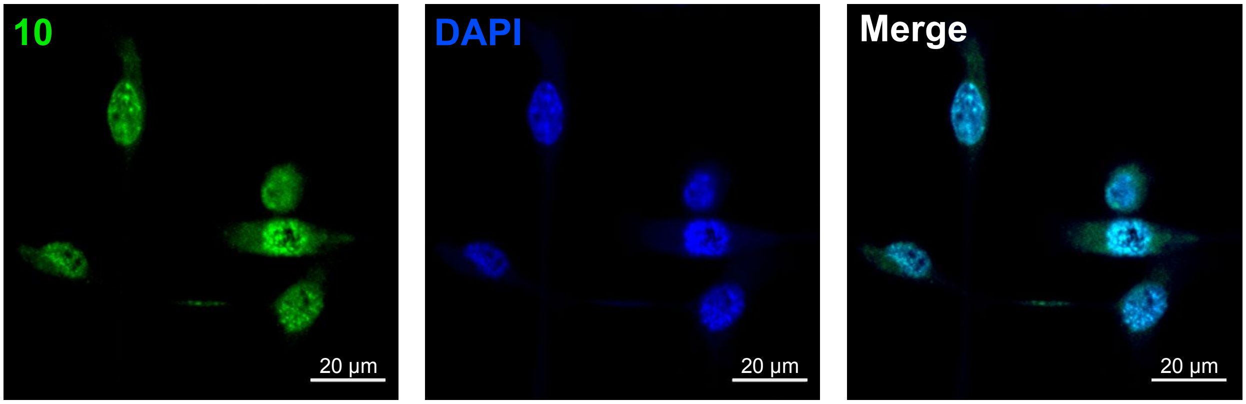

**Figure S8. Compound 10 localizes to the nuclei.** The compound **10** (green) (NucShine) localizes to the nuclei (blue) in BV2 microglia. Scale bars represent 20 µm.

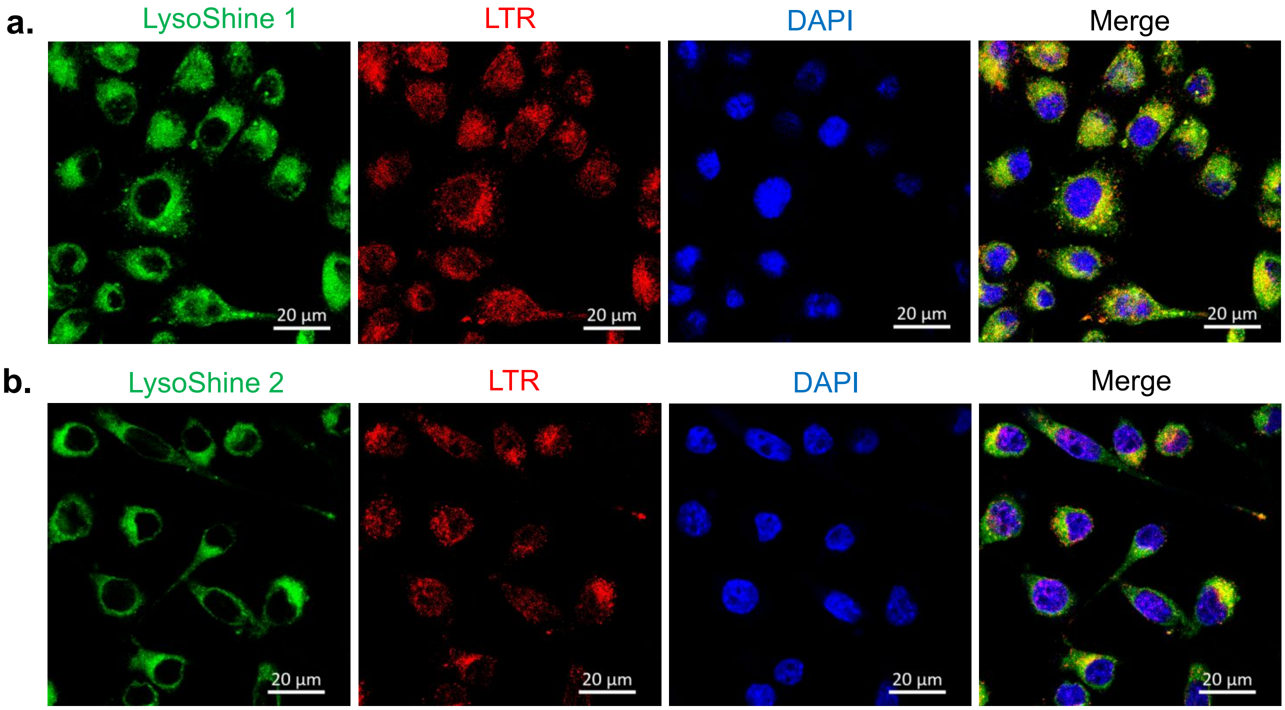

**Figure S9.** Confocal Imaging of Lysosomal probes LysoShine 1 and LysoShine 2 in BV2 microglia. Confocal images depicting the co-colocalization of (**a**) LysoShine 1 and (**b**) LysoShine 2 lysosomal probes (green)

with LysoTracker Red DND-99 (LTR, red) in BV2 microglia. Nuclei are stained with DAPI (blue). Scale bars represent 20 μm.

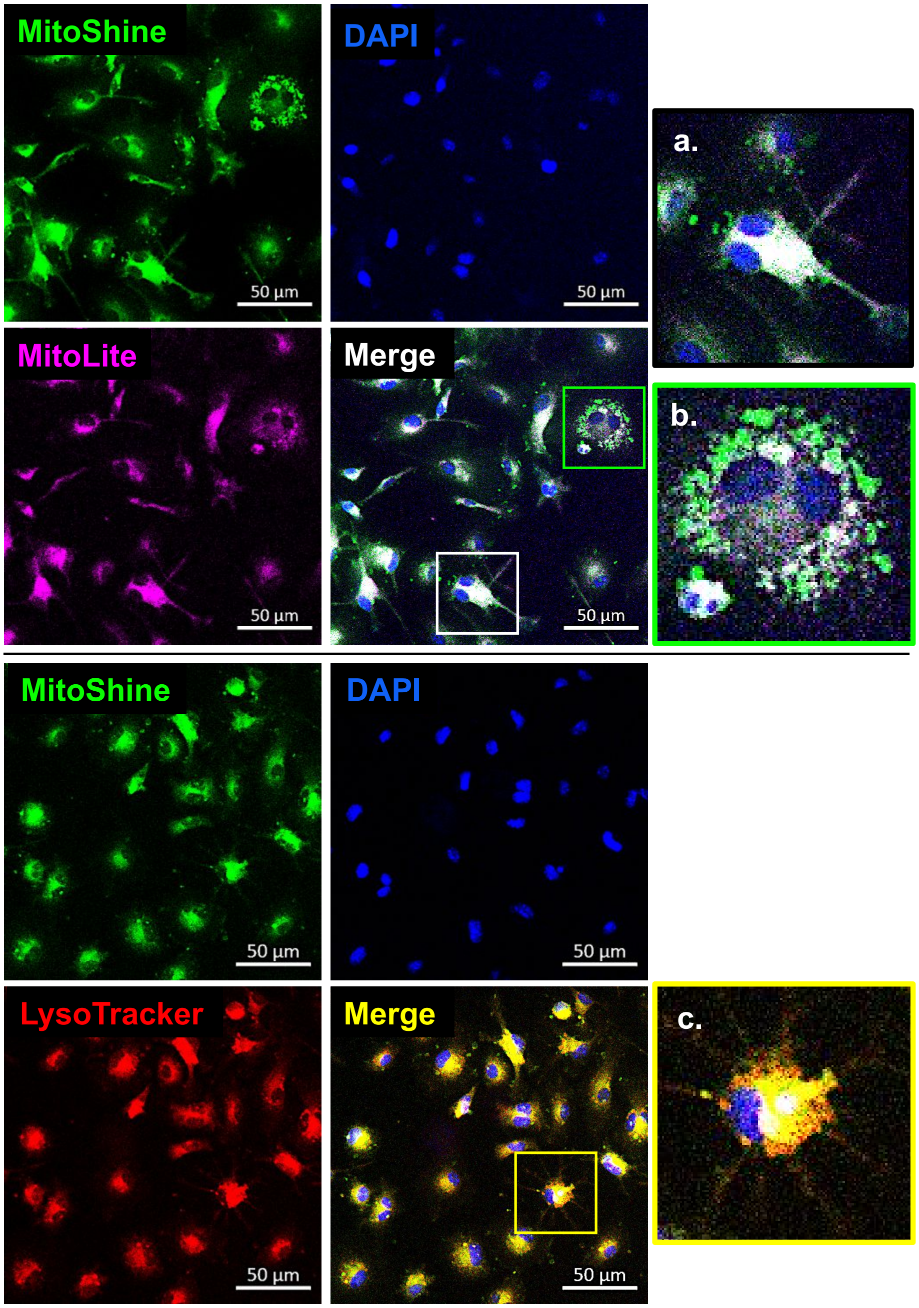

**Figure S10. Confocal imaging of primary mouse microglial cells with MitoShine.** The localization of the compound was observed in mitochondria (magenta) and acidic lysosomal organelles (red). Magnified images on the far-right show MitoShine localization in (**a**) non-mitochondria (likely lysosomes), (**b**) mitochondria, and (**c**) lysosomes. Nuclear DNA is stained with DAPI (blue). Scale bars represent 50 μm.

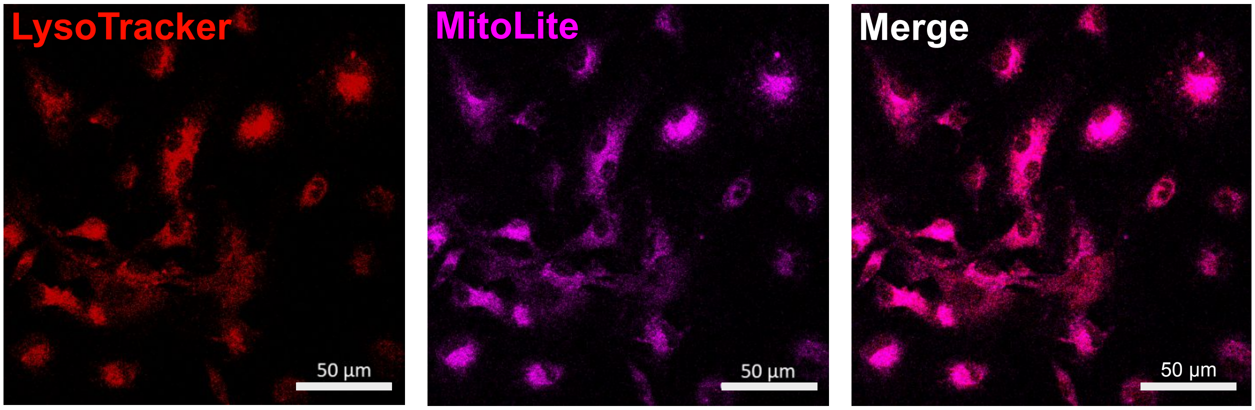

**Figure S11.** Overlap of lysosomes and mitochondria in primary microglia. The overlap of mitochondria (magenta) and acidic lysosomal organelles (red) observed via confocal microscope during co-treatment with MitoShine florescent probe. Scale bars represent 50 µm.

### EXPERIMENTAL SECTION

Unless noted otherwise, all reagents and solvents were purchased from commercial sources and used as received. All reactions were performed in either round bottom flask with septum or microwave vial with seal septum. The proton ($^1H$) and carbon ($^{13}C$) NMR spectra were obtained using a 500 MHz using $Me_4Si$ as an internal standard and are reported in $\delta$ units. Coupling constants (*J* values) are reported in Hz. Silica gel column chromatography was performed on Teledyne ISCO (EZprep model) instrument. High-resolution mass spectra (HRMS) were obtained using the electron spray ionization (ESI) technique and as TOF mass analyzer. Organic solvents and starting materials were used as received. The absorption and fluorescence spectra were process by GraphPad Prism software (version 9)

#### Synthetic Procedures

**Procedure for the synthesis of Diethyl 10-(4-(dimethylamino)phenyl)-5,5-difluoro-1,3,7,9-tetramethyl-5*H*-4λ$^4$,5λ$^4$-dipyrrolo[1,2-c:2',1'-*f*][1,3,2]diazaborinine-2,8-dicarboxylate (5, kpgc02s254)**

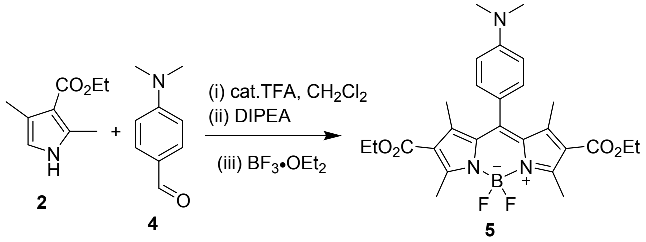

A clean oven dried 500 mL round bottom flask with a magnetic stir bar, charged with 2,4-Dimethyl-1H-pyrrole-3-carboxylic acid ethyl ester (2 equiv., 3 mmol, 502 mg) and 4-(Dimethylamino) benzaldehyde (1 equiv., 1.5 mmol, 224 mg) in anhydrous $CH_2Cl_2$ (300 mL). The solution was purged with Argon twice. Then, 1 drop of trifluoro acetic acid was added under inert reaction condition and reaction was allowed to stir at room temperature for 24 hours. The reaction was monitored by TLC. Next, septum was removed and DDQ (1 equiv., 1.5 mmol) was added quickly. Again, the reaction mixture was purged with Argon and stirred at room temperature for 15 min. The immediate color change to dark purple was observed. The solvent was partially removed and compound was isolated over short pad of alumina (neutral) using 1-10% methanol in dichloromethane as an eluent as a dark red solid powder. The product was immediately used for the next step.

In a clean oven dried round bottom flask (500 mL) with a stir bar, a mixture of the resulting compound, anhydrous DCM (200 mL) and diisopropyl ethylamine (5 mL) was added under inert atmosphere. The solution was stirred for 10 minutes. $BF_3$•$OEt_2$ (5 mL) was added slowly and stirred for additional 5 hours under inert atmosphere. The crude product was extracted with dichloromethane: water, washed with brine, dried over sodium sulfate. Further, compound was purified using flash silica column chromatography with 0-1% methanol in dichloromethane as an eluent and a brownish-purple solid product (460 mg, yield 60%) was obtained. If needed, the purification can be repeated. $^1H$ NMR (500 MHz, $CDCl_3$): $\delta$ 7.03 (d, *J* = 8.8 Hz, 2H), 6.80 (d, *J* = 8.7 Hz, 2H), 4.28 (q, *J* = 7.1, 7.1, 7.1 Hz, 4H), 3.05 (s, 6H), 2.82 (d, *J* = 1.4 Hz, 6H), 1.78 (s, 6H), 1.33 (t, *J* = 7.1, 7.1 Hz, 6H); $^{13}C$ NMR (126 MHz, $CDCl_3$) δ 190.36, 164.52, 158.76, 151.15, 147.76, 147.60,

132.20, 128.80, 122.13, 121.14, 112.43, 111.01, 60.13, 40.23, 40.11, 14.96, 14.32, 14.16; HRMS (ESI) m/z: [M - H]$^+$ calcd for $C_{27}H_{31}BF_2N_3O_4$ 510.2376; Found 510.2380.

**Procedure for the synthesis of diethyl (*E*)-10-(4-(dimethylamino)phenyl)-5,5-difluoro-3-(4-hydroxystyryl)-1,7,9-trimethyl-5*H*-4l4,5l4-dipyrrolo[1,2-c:2',1'-*f*][1,3,2]diazaborinine-2,8-dicarboxylate (6a, kpgc02s264)**

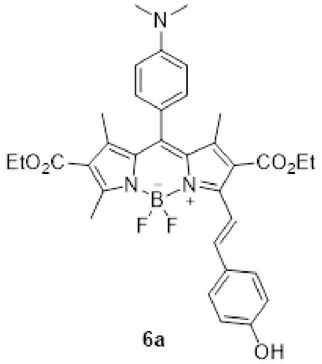

6a

In a clean oven dried microwave vial with a stir bar, a mixture of compound **5** (30 mg, 0.058 mmol, 1 equiv), 4-hydroxybenzaldehyde (7 mg, 0.058 mmol, 1 equiv), piperidine (100 uL), acetic acid (100 uL), activated molecular sieve (500 mg) were added and the vial was sealed and purged with Argon. Then, anhydrous toluene (2 mL) was added, and reaction mixture was stirred at 120 °C (reflux) for 45 minutes. The reaction was monitored by TLC (Silica, 10% ethylacetate in dichloromethane). The reaction mixture was cooled to room temperature and washed three times with water. The organic phase was dried over sodium sulfate and the organic solvent was evaporated under reduced pressure. The residue was purified by silica gel flash column chromatography (CombiFlash) using 0-20% ethylacetate in dichloromethane to afford the desired compound **6a** as reddish-purple solid (13 mg, yield 37%). $^1$H NMR (500 MHz, Acetone) δ 8.58 (s, 1H), 7.25 – 7.19 (m, 2H), 6.90 (dd, *J* = 8.7, 4.9 Hz, 4H), 6.71 (d, *J* = 8.6 Hz, 2H), 6.55 (d, *J* = 16.3 Hz, 1H), 5.87 (d, *J* = 16.2 Hz, 1H), 4.25 (dq, *J* = 19.4, 7.1, 7.1, 7.1 Hz, 4H), 3.05 (s, 6H), 2.69 (s, 3H), 1.86 (s, 3H), 1.32 (t, *J* = 7.1, 7.1 Hz, 3H), 1.19 (t, *J* = 7.1, 7.1 Hz, 3H); $^{13}$C NMR (126 MHz, $CDCl_3$) *δ* 212.05, 169.69, 168.64, 163.48, 162.65, 161.54, 156.34, 152.20, 151.45, 148.78, 141.03, 136.87, 136.71, 134.73, 133.47, 132.67, 127.02, 125.27, 125.08, 121.64, 120.46, 117.71, 65.61, 65.25, 19.76, 19.25, 19.20, 19.13, 18.83; HRMS (ESI) m/z: [M + H]$^+$ calcd for $C_{34}H_{37}BF_2N_3O_5$ 616.2794; Found 616.2799.

**Procedure for the synthesis of diethyl (*E*)-10-(4-(Dimethylamino)phenyl)-5,5-difluoro-3-(4-hydroxy-3-nitrostyryl)-1,7,9-trimethyl-5*H*-4l4,5l4-dipyrrolo[1,2-c:2',1'-*f*][1,3,2]diazaborinine-2,8-dicarboxylate (6b, kpgc02s270)**

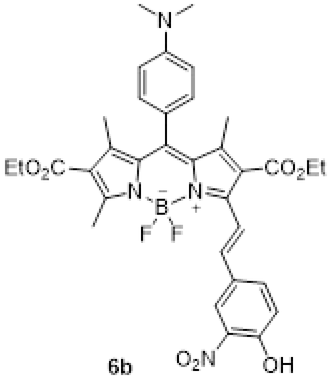

6b

In a clean oven dried microwave vial with a stir bar, a mixture of compound **5** (51.1 mg, 0.1 mmol, 1 equiv), 4-hydroxy-3-nitro benzaldehyde (16.7 mg, 0.1 mmol, 1 equiv), piperidine (100 uL), acetic acid (100 uL), activated molecular sieve (500 mg) were added and purged with Argon. Then, anhydrous toluene (2 mL) was added, and reaction mixture was stirred at 120 °C (reflux) for 12 hours. The reaction was monitored by TLC (Silica, 10% ethylacetate in dichloromethane). The reaction mixture was cooled to room temperature and washed three times with water. The organic phase was dried over sodium sulfate and the organic solvent was evaporated under reduced pressure. The residue was purified by silica gel flash column chromatography (CombiFlash) using 0-30% ethylacetate in dichloromethane to afford the desired compound **6a** as dark purple solid (29 mg, yield 45%); $^1$H NMR (500 MHz, $CDCl_3$) δ 10.73 (s, 1H), 8.48 – 8.42 (m, 2H), 8.22 (d, *J* = 2.2 Hz, 1H), 7.92 (dd, *J* = 8.8, 2.2 Hz, 1H), 7.66 – 7.54 (m, 3H), 7.40 (d, *J* = 16.5 Hz, 1H), 7.22 (d, *J* = 8.8 Hz, 1H), 4.32 (dq, *J* = 18.1, 7.1, 7.1, 7.1 Hz, 4H), 2.88 (s, 3H), 1.67 (s, 3H), 1.54 (s, 6H), 1.32 (dt, *J* = 14.6, 7.1, 7.1 Hz, 6H); $^{13}$C NMR (126 MHz, $CDCl_3$) δ 164.95, 163.85, 161.44, 155.68, 152.31, 148.79, 147.28, 144.16, 141.15, 137.16, 135.39, 133.66,

131.64, 129.65, 124.84, 124.47, 120.76, 118.20, 61.29, 60.58, 29.72, 15.31, 14.23, 14.11, 13.77; HRMS (ESI) m/z: $[M + H]^+$ calcd for $C_{34}H_{36}BF_2N_4O_7$ 661.2645; Found 661.2652.

**Procedure for the synthesis of Diethyl 5,5-difluoro-1,3,7,9-tetramethyl-10-(4-nitrophenyl)-5*H*-4$\lambda^4$,5$\lambda^4$-dipyrrolo[1,2-c:2',1'-*f*][1,3,2]diazaborinine-2,8-dicarboxylate (8, kpgc02s273)**

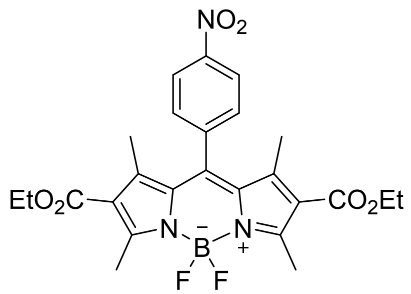

4-Nitrobenzaldehyde (2.0 mmol, 302 mg, 1 equiv) and 2,4-Dimethyl-1*H*-pyrrole-3-carboxylic acid ethyl ester (4.0 mmol, 669 mg, 2 equiv) were dissolved in 350 mL of anhydrous $CH_2Cl_2$ under Argon atmosphere. One drop of TFA was added, and the solution was stirred at room temperature overnight. When TLC monitoring (silica; $CH_2Cl_2$) showed complete consumption of the aldehyde, a solution of 2,3-Dichloro-5,6-dicyano-1,4-benzoquinone (DDQ, 908 mg, 4.0 mmol) in $CH_2Cl_2$ was added, and stirring was continued for 20 minutes under Argon atmosphere. The reaction mixture was washed with water, dried over sodium sulfate, filtered, and evaporated. The compound was purified by short (manual) column chromatography over neutral alumina ($CH_2Cl_2$). The brown powder thus obtained and 5 mL of *N,N*-Diisopropylethylamine (DIPEA) were dissolved in 200 mL of toluene under an Argon atmosphere. Then 5 mL of $BF_3{\bullet}Et_2O$ was added dropwise, and the solution was stirred at room temperature for 30 min. The reaction mixture was washed with water, dried over sodium sulfate, filtered, and evaporated. The compound was purified by silica gel column chromatography ($CH_2Cl_2$/n-hexane = 1/1) to give an orange-red solid (620 mg, yield 60%). $^1H$ NMR (500 MHz, $CDCl_3$) δ 8.44 (d, *J* = 8.7 Hz, 2H), 7.57 – 7.52 (m, 2H), 4.29 (q, *J* = 7.2, 7.1, 7.1 Hz, 4H), 2.84 (s, 6H), 1.64 (s, 6H), 1.33 (t, *J* = 7.1, 7.1 Hz, 6H); $^{13}C$ NMR (126 MHz, $CDCl_3$) δ 163.96, 160.60, 148.74, 146.97, 142.28, 141.22, 130.72, 129.52, 124.82, 123.14, 60.48, 15.13, 14.27, 14.02; HRMS (ESI) m/z: $[M - H]^+$ calcd for $C_{25}H_{25}BF_2N_3O_6$ 512.1805; Found 512.1801.

**Procedure[1] for the synthesis of diethyl 10-(4-aminophenyl)-5,5-difluoro-1,3,7,9-tetramethyl-5*H*-4l4,5l4-dipyrrolo[1,2-c:2',1'-*f*][1,3,2]diazaborinine-2,8-dicarboxylate (9, kpgc02s274)**

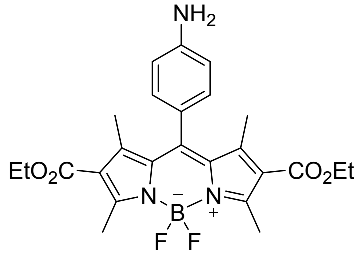

To a solution of nitro-compound **8** (0.5 mmol) in 120 mL of degassed EtOH:$CH_2Cl_2$ (1:1), was added a suspension of Pd/C (50 mg, 10% mol) in EtOH under inert atmosphere. The resulting mixture was stirred at room temperature under $H_2$ (1 atm, balloon) for 12 hours. When the reaction was completed, inorganic solids were removed by filtration through Celite® pad and washed with several portions of $CH_2Cl_2$. The TLC showed a small amount of impurities. The product was purified by flash chromatography using 0-20% $CH_2Cl_2$: MeOH. The silica in cartridge and column was neutralized by passing acetone (containing 0.5% triethylamine). The orange colored solid was obtained quantitatively. $^1H$ NMR (500 MHz, $CDCl_3$) δ 6.99 (d, *J* = 8.4 Hz, 2H), 6.81 (d, *J* = 8.4 Hz, 2H), 4.28 (q, *J* = 7.1, 7.1, 7.1 Hz, 4H), 2.82 (s, 6H), 1.78 (s, 6H), 1.33 (t, *J* = 7.1, 7.1 Hz, 6H); $^{13}C$ NMR (126 MHz, $CDCl_3$) δ 164.44, 159.01, 147.82, 146.95, 132.05, 128.91, 123.81, 122.29, 115.69, 60.17, 14.96, 14.31, 14.05; HRMS (ESI) m/z: $[M + H]^+$ calcd for $C_{25}H_{29}BF_2N_3O_4$ 484.2219; Found 484.2225.

**Procedure for the synthesis of diethyl 10-(4-(2-bromoacetamido)phenyl)-5,5-difluoro-1,3,7,9-tetramethyl-5*H*-4l4,5l4-dipyrrolo[1,2-c:2',1'-*f*][1,3,2]diazaborinine-2,8-dicarboxylate (10, kpgc02s276)**

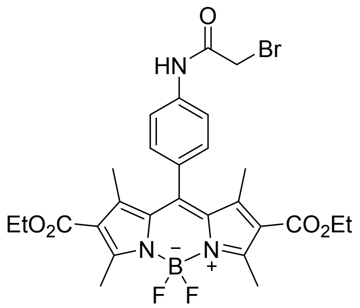

A solution of $NH_2$-compound **9** (60 mg, 0.12 mmol) in $CH_2Cl_2$ (3 mL) was kept in ice-bath. Triethylamine (0.14 mmol, 19 µL) was added to a solution and then bromoacetyl bromide (0.14 mmol, 12 µL) was added slowly. It was stirred at room temperature. Once precipitation was observed within 5 minutes, reaction mixture was directly quenched with 10 mL $CH_2Cl_2$ and saturated sodium bicarbonate solution. The organic layer was collected and removed under reduced pressure. The product was obtained in quantitative yield as bright orange solid (72 mg, 97% yield). $^1$H NMR (500 MHz, $CDCl_3$): *δ* 8.48 (bs, 1H), 7.82 – 7.76 (m, 2H), 7.29 – 7.21 (m, 2H), 4.27 (q, *J* = 7.1, 7.1, 7.1 Hz, 4H), 4.06 (s, 2H), 2.82 (s, 6H), 1.70 (s, 6H), 1.32 (t, *J* = 7.1, 7.1 Hz, 6H); $^{13}$C NMR (126 MHz, $CDCl_3$): *δ* 164.28, 163.73, 159.61, 147.57, 145.06, 138.64, 131.56, 130.68, 128.73, 122.59, 120.49, 60.31, 46.23, 29.39, 15.03, 14.28, 13.95; HRMS (ESI) m/z: $[M + H]^+$ calcd for $C_{27}H_{30}BBrF_2N_3O_5$ 604.14030; Found 604.1438.

**Procedure for the synthesis of Diethyl 10-(4-(2-(diethylamino)acetamido)phenyl)-5,5-difluoro-1,3,7,9-tetramethyl-5*H*-4λ$^4$,5λ$^4$-dipyrrolo[1,2-c:2',1'-*f*][1,3,2]diazaborinine-2,8-dicarboxylate (LysoShine 1, kpgc02s277)**

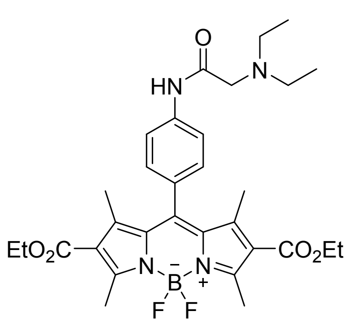

In a screw cap vial, to a solution of **10** (20 mg, 0.033 mmol) in acetone (1 mL) was added diethylamine (0.066 mmol, 2 equiv, 7 µL) and stirred at 60 °C for 1 hour. The reaction was monitored by TLC (silica, $CH_2Cl_2$: EtOAc 9:1). Immediately, the reaction mixture was directly loaded on a cartridge and product was eluted in $CH_2Cl_2$: MeOH (95:5) using flash chromatography. The product was obtained as orange solid (11 mg, 56% yield). $^1$H NMR (500 MHz, $CDCl_3$): δ 9.63 (bs, 1H), 7.79 (d, *J* = 8.5 Hz, 2H), 7.22 (d, *J* = 8.5 Hz, 2H), 4.27 (q, *J* = 7.1, 7.1, 7.1 Hz, 4H), 3.19 (s, 2H), 2.82 (s, 6H), 2.69 (q, *J* = 7.1, 7.1, 7.1 Hz, 4H), 2.16 (s, 1H), 1.72 (s, 6H), 1.32 (t, *J* = 7.2, 7.2 Hz, 6H), 1.13 (t, *J* = 7.1, 7.1 Hz, 6H); $^{13}$C NMR (126 MHz, $CDCl_3$) δ 170.50, 164.30, 159.46, 147.65, 145.60, 139.19, 131.67, 129.58, 128.64, 122.51, 119.89, 60.24, 58.13, 48.75, 14.99, 14.29, 14.03, 12.41; HRMS (ESI) m/z: $[M + H]^+$ calcd for $C_{31}H_{40}BF_2N_4O_5$ 597.3060; Found 597.3065.

**Procedure for the synthesis of diethyl 5,5-difluoro-1,3,7,9-tetramethyl-10-(4-(2-((2-morpholinoethyl)amino)acetamido)phenyl)-5*H*-4λ$^4$,5λ$^4$-dipyrrolo[1,2-c:2',1'-*f*][1,3,2]diaza-borinine-2,8-dicarboxylate (LysoShine 2, kpgc02s280)**

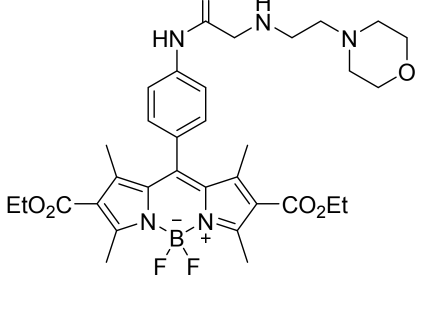

To a solution of **10** compound (20 mg, 0.033 mmol) in acetone (1 mL) was added 4-(2-aminoethyl)morpholine (0.066 mmol, 2 equiv, ~9 µL) and stirred at 60 °C for 1 hour. The reaction was monitored by TLC (silica, $CH_2Cl_2$: EtOAc 9:1) and a new polar product was observed. Immediately, the reaction mixture was directly loaded on a cartridge and product was eluted with gradient of 0-10% methanol in dichloromethane. The product was obtained as Dark orange solid (10 mg, 46% yield). $^1H$ NMR (500 MHz, $CDCl_3$) δ 9.68 (bs, 1H), 7.82 (d, *J* = 8.5 Hz, 2H), 7.22 (d, *J* = 8.5 Hz, 2H), 4.27 (q, *J* = 7.1, 7.1, 7.1 Hz, 4H), 3.71 (t, *J* = 4.7, 4.7 Hz, 4H), 3.43 (s, 2H), 2.82 (s, 8H), 2.56 – 2.50 (m, 2H), 2.50 – 2.44 (m, 4H), 1.71 (s, 6H), 1.32 (t, *J* = 7.1, 7.1 Hz, 6H); $^{13}C$ NMR (126 MHz, $CDCl_3$) δ 170.44, 164.30, 159.46, 147.63, 145.55, 139.21, 131.65, 129.63, 128.60, 122.51, 120.06, 66.94, 60.26, 58.11, 53.71, 53.18, 46.39, 15.01, 14.29, 13.95; HRMS (ESI) m/z: $[M + H]^+$ calcd for $C_{33}H_{43}BF_2N_5O_6$ 654.3274; Found 654.3280.

**Procedure for the synthesis of (2-((4-(2,8-bis(ethoxycarbonyl)-5,5-difluoro-1,3,7,9-tetramethyl-5*H*-4l4,5l4-dipyrrolo[1,2-c:2',1'-*f*][1,3,2]diazaborinin-10-yl)phenyl)amino)-2-oxoethyl)triphenylphosphonium bromide (MitoShine, kpgc02s286)**

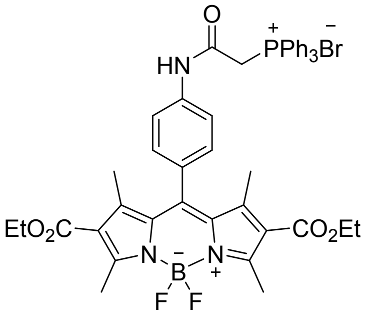

To a solution of **10** (37 mg, 0.06 mmol) in anhydrous acetonitrile (2 mL), triphenylphoshine (0.18 mmol, 3 equiv, 48 mg) was added and stirred at reflux condition overnight. The reaction was monitored by TLC (silica, $CH_2Cl_2$: EtOAc 9:1) and a new polar product was observed. Immediately, the reaction mixture was directly loaded on a cartridge and product was eluted in $CH_2Cl_2$: MeOH (90:10). The product was obtained as greenish orange solid (30 mg, 63% yield). $^1H$ NMR (500 MHz, $CDCl_3$) δ 11.68 (bs, 1H), 7.90 – 7.76 (m, 10H), 7.70 – 7.63 (m, 6H), 7.14 – 7.09 (m, 2H), 5.16 (d, *J* = 14.4 Hz, 2H), 4.28 (q, *J* = 7.1, 7.1, 7.1 Hz, 4H), 2.81 (s, 6H), 2.63 (s, 1H), 2.17 (s, 2H), 1.65 (s, 6H), 1.36 – 1.30 (m, 6H); $^{13}C$ NMR (126 MHz, $CDCl_3$) δ 164.34, 147.73, 135.27, 134.11, 134.03, 130.34, 130.23, 128.17, 120.83, 117.64, 60.23, 29.28, 14.98, 14.29, 13.85; HRMS (ESI) m/z: $[M - Br]^+$ calcd for $C_{45}H_{44}BF_2N_3O_5P^+$ 786.3074; Found 786.3079.

#### $^{1}H$ and $^{13}C$ NMR spectra

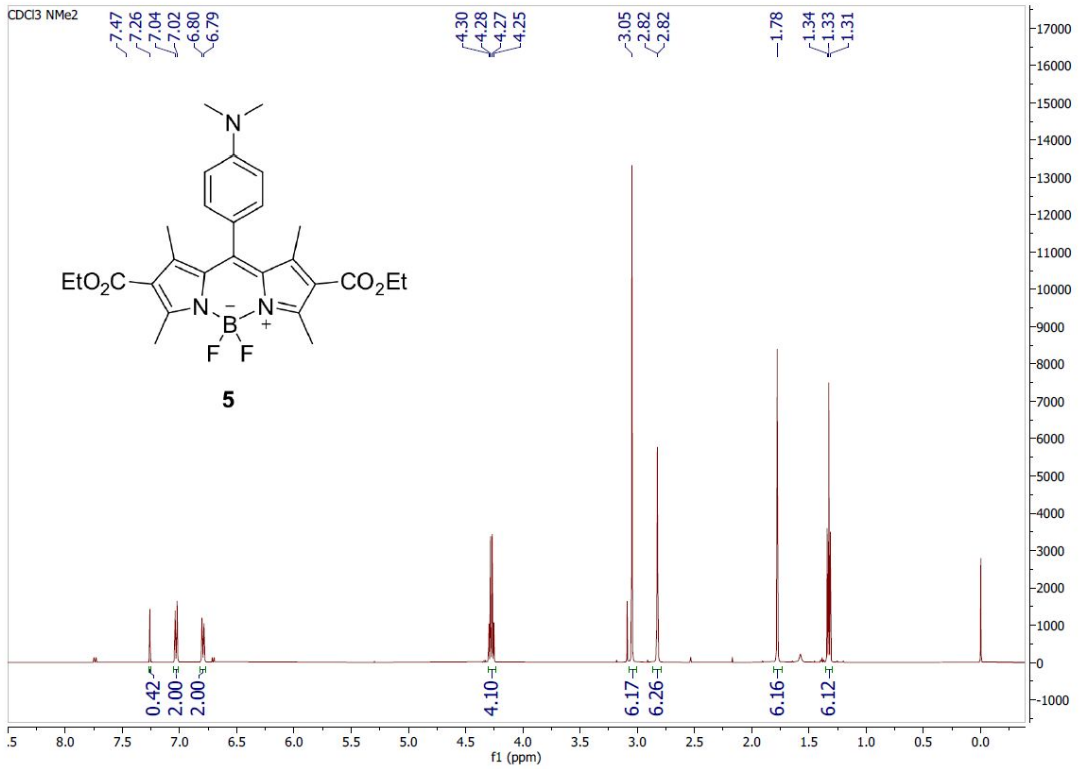

**$^{1}$H-spectrum of Diethyl 10-(4-(dimethylamino)phenyl)-5,5-difluoro-1,3,7,9-tetramethyl-5*H*-4λ$^{4}$,5λ$^{4}$-dipyrrolo[1,2-c:2',1'-*f*][1,3,2]diazaborinine-2,8-dicarboxylate (5)**

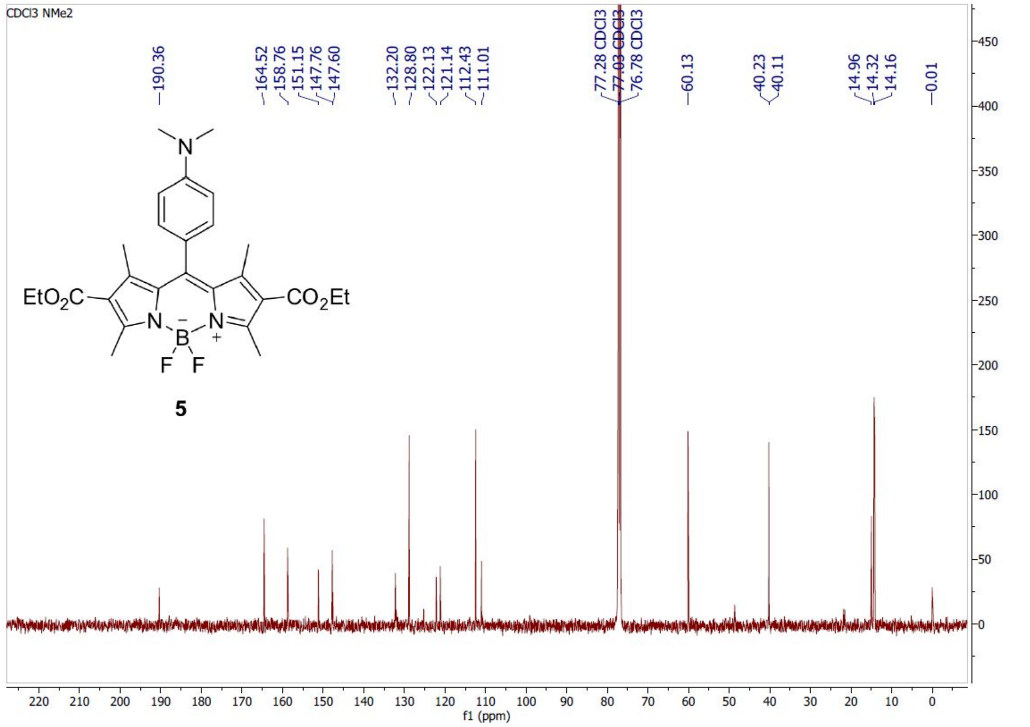

**$^{13}$C-spectrum of Diethyl 10-(4-(dimethylamino)phenyl)-5,5-difluoro-1,3,7,9-tetramethyl-5*H*-4λ$^{4}$,5λ$^{4}$-dipyrrolo[1,2-c:2',1'-*f*][1,3,2]diazaborinine-2,8-dicarboxylate (5)**

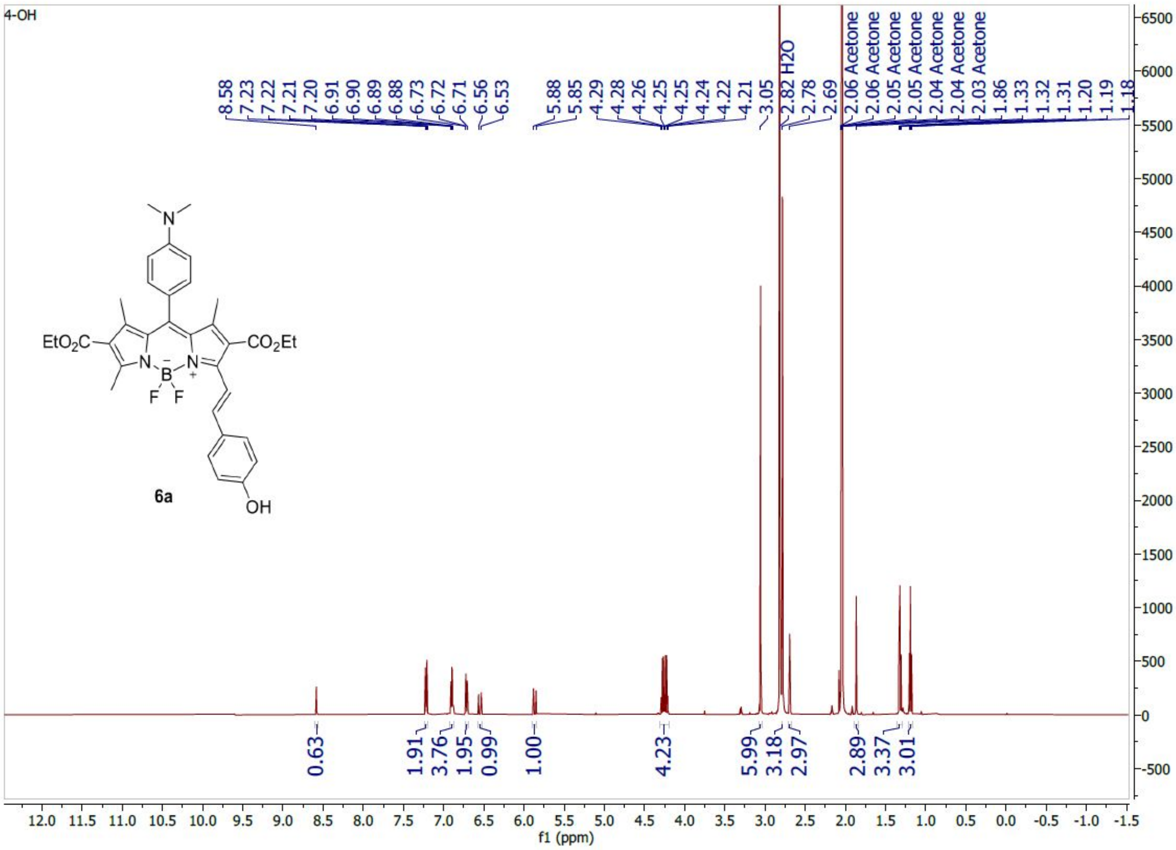

[1]H-spectrum of (*E*)-10-(4-(Dimethylamino)phenyl)-5,5-difluoro-3-(4-hydroxystyryl)-1,7,9-trimethyl-5*H*-4l4,5l4-dipyrrolo[1,2-c:2',1'-*f*][1,3,2]diazaborinine-2,8-dicarboxylate (6a)

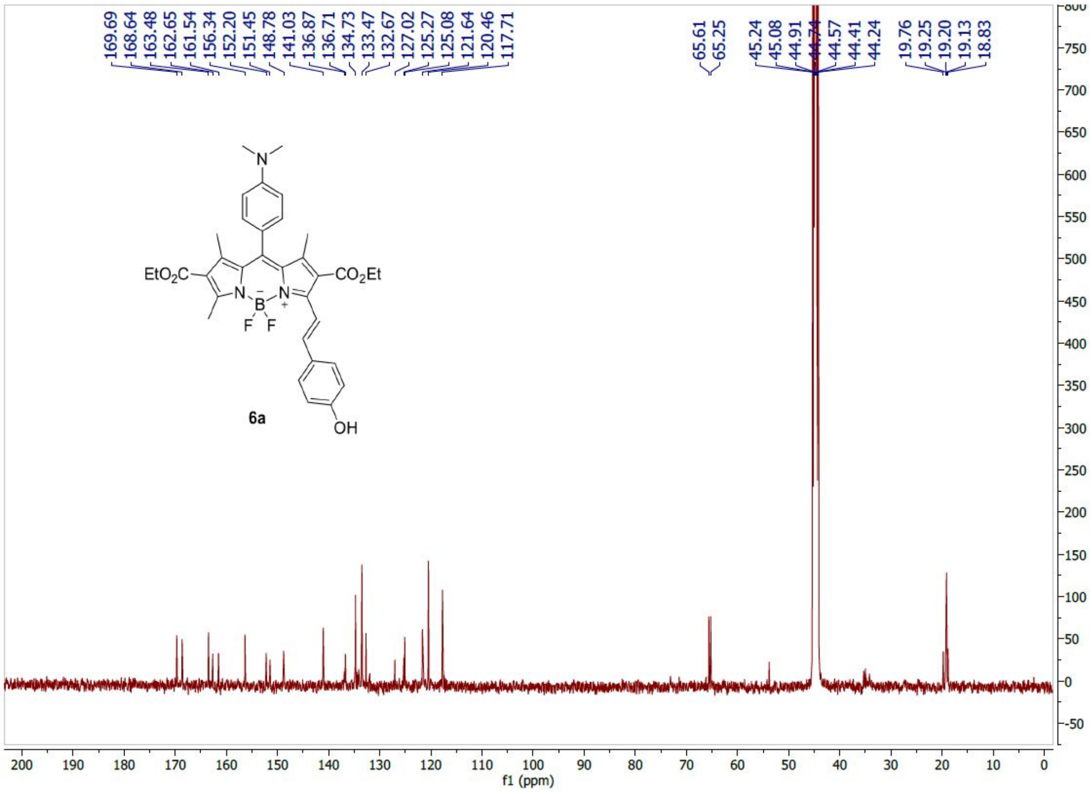

[13]C-spectrum of (*E*)-10-(4-(Dimethylamino)phenyl)-5,5-difluoro-3-(4-hydroxystyryl)-1,7,9-trimethyl-5*H*-4l4,5l4-dipyrrolo[1,2-c:2',1'-*f*][1,3,2]diazaborinine-2,8-dicarboxylate (6a)

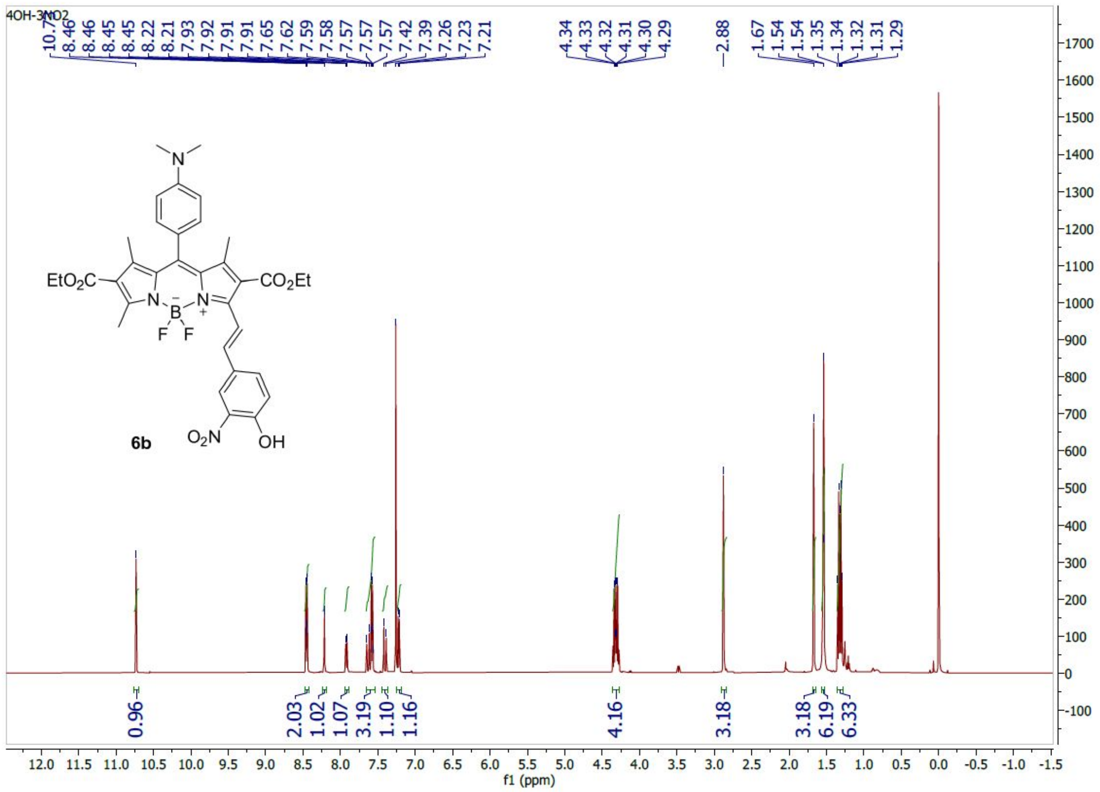

**[1]H-spectrum Diethyl (*E*)-10-(4-(dimethylamino)phenyl)-5,5-difluoro-3-(4-hydroxy-3-nitrostyryl)-1,7,9-trimethyl-5*H*-4l4,5l4-dipyrrolo[1,2-c:2',1'-*f*][1,3,2]diazaborinine-2,8-dicarboxylate (6b)**

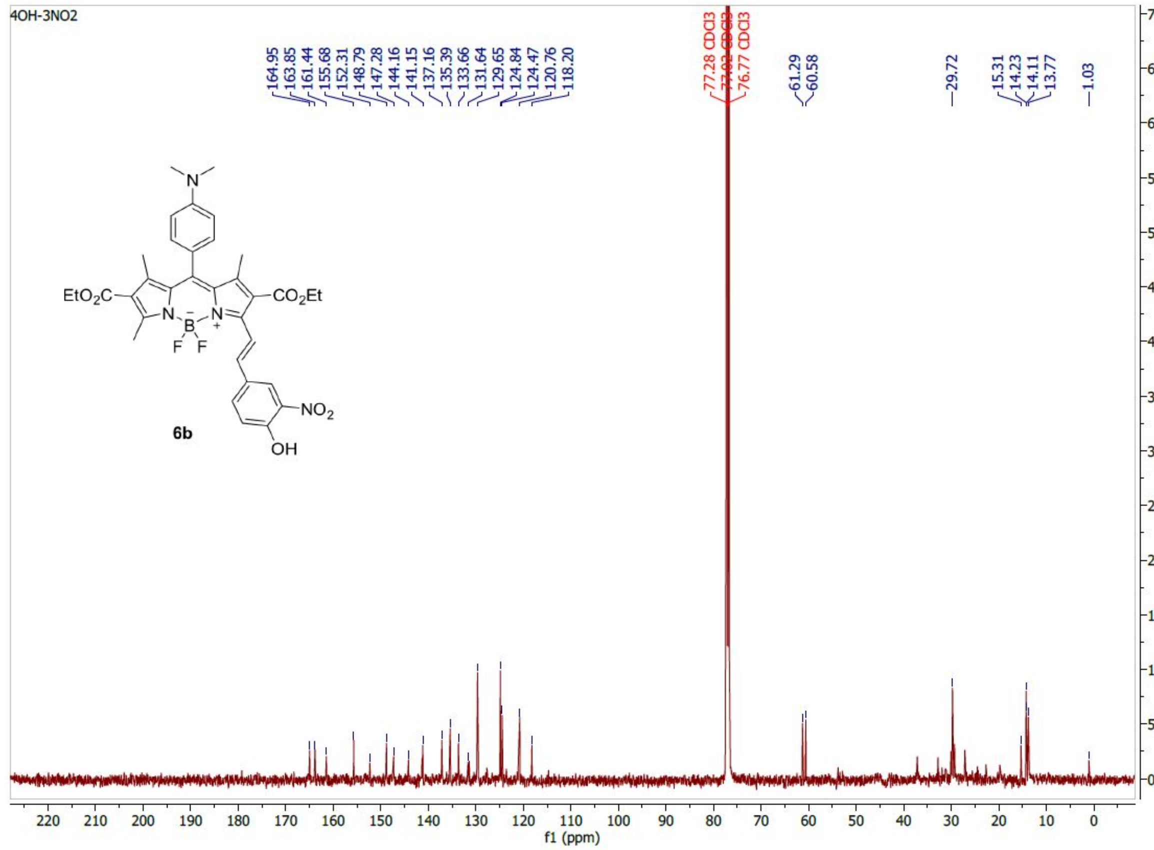

**[13]C-spectrum of Diethyl (*E*)-10-(4-(dimethylamino)phenyl)-5,5-difluoro-3-(4-hydroxy-3-nitrostyryl)-1,7,9-trimethyl-5*H*-4l4,5l4-dipyrrolo[1,2-c:2',1'-*f*][1,3,2]diazaborinine-2,8-dicarboxylate (6b)**

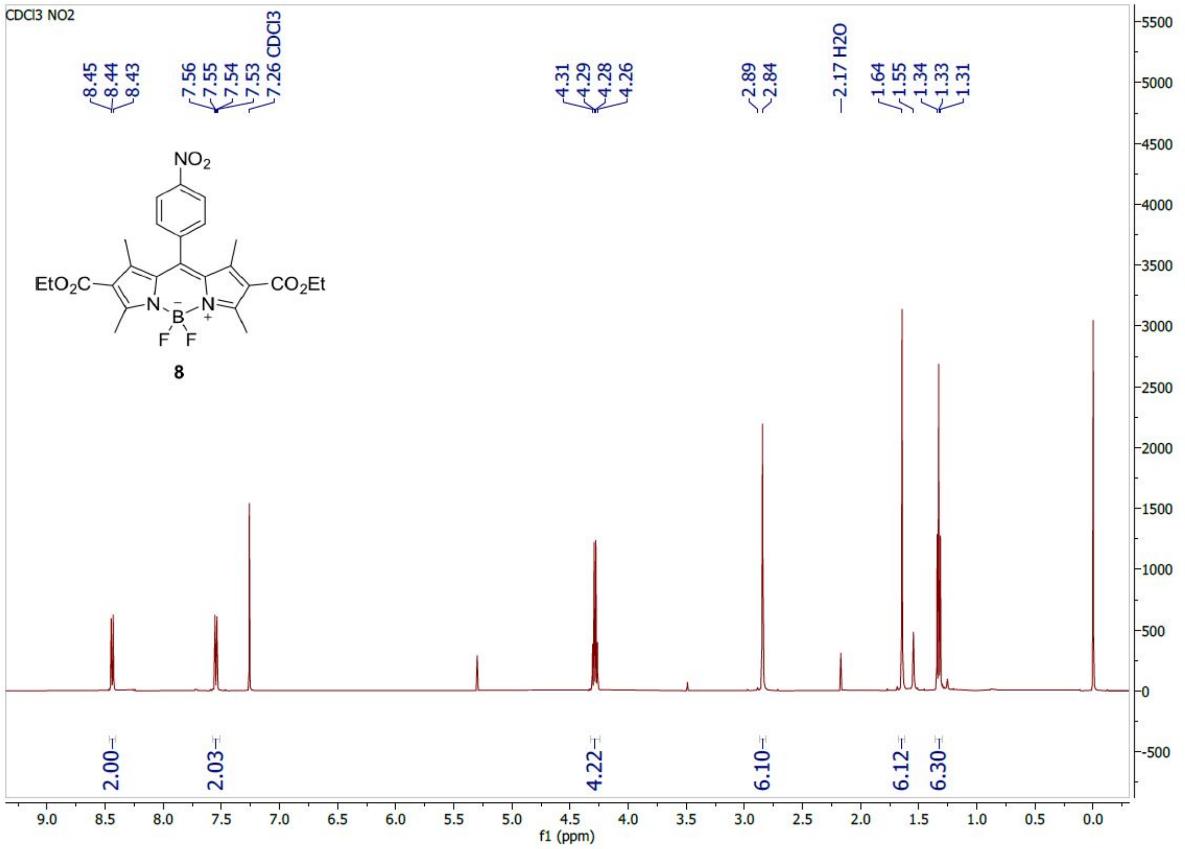

**$^1$H-spectrum of Diethyl 5,5-difluoro-1,3,7,9-tetramethyl-10-(4-nitrophenyl)-5*H*-4λ$^4$,5λ$^4$-dipyrrolo[1,2-c:2',1'-*f*][1,3,2]diazaborinine-2,8-dicarboxylate (**8**)**

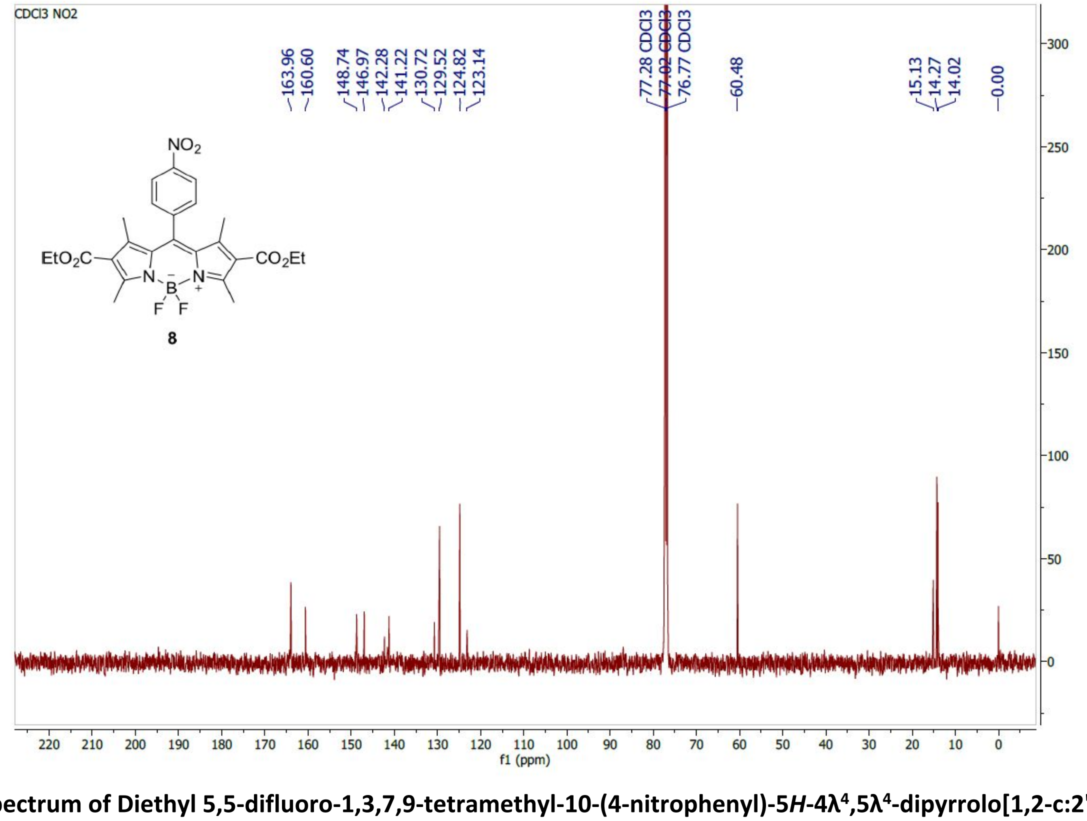

**$^{13}$C-spectrum of Diethyl 5,5-difluoro-1,3,7,9-tetramethyl-10-(4-nitrophenyl)-5*H*-4λ$^4$,5λ$^4$-dipyrrolo[1,2-c:2',1'-*f*][1,3,2]diazaborinine-2,8-dicarboxylate (**8**)**

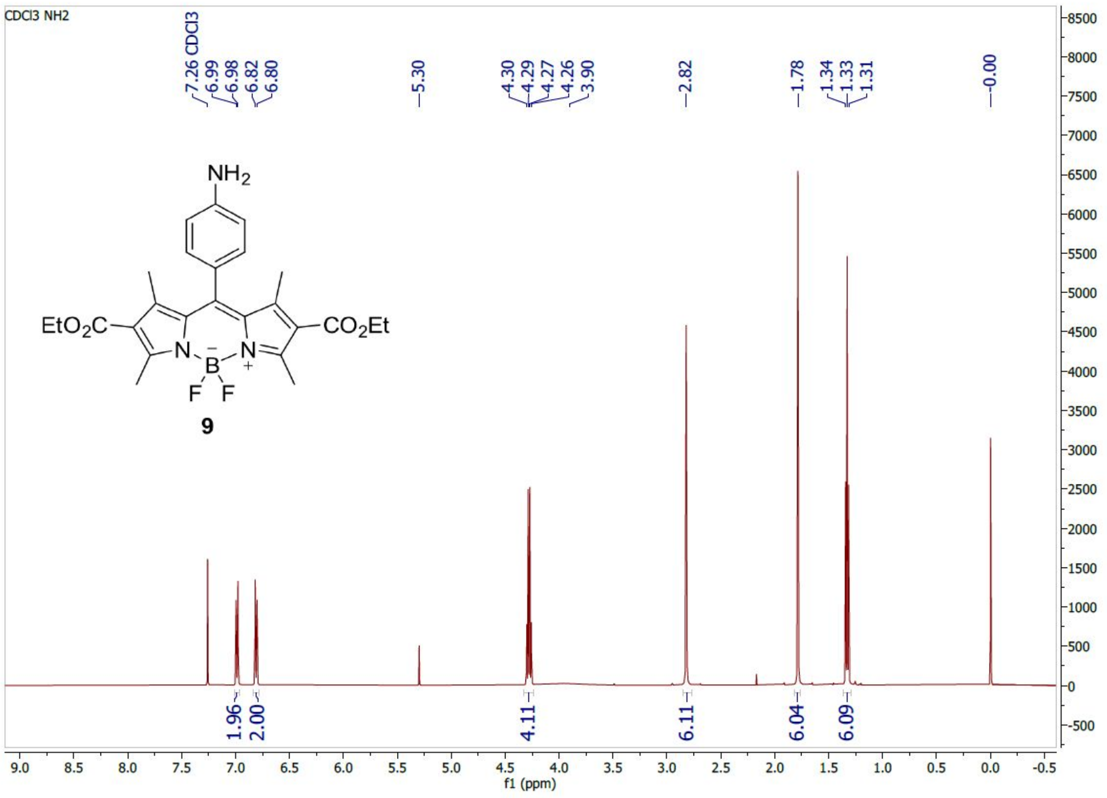

[1]H-spectrum of Diethyl 10-(4-aminophenyl)-5,5-difluoro-1,3,7,9-tetramethyl-5*H*-4l4,5l4-dipyrrolo[1,2-c:2',1'-*f*][1,3,2]diazaborinine-2,8-dicarboxylate (9)

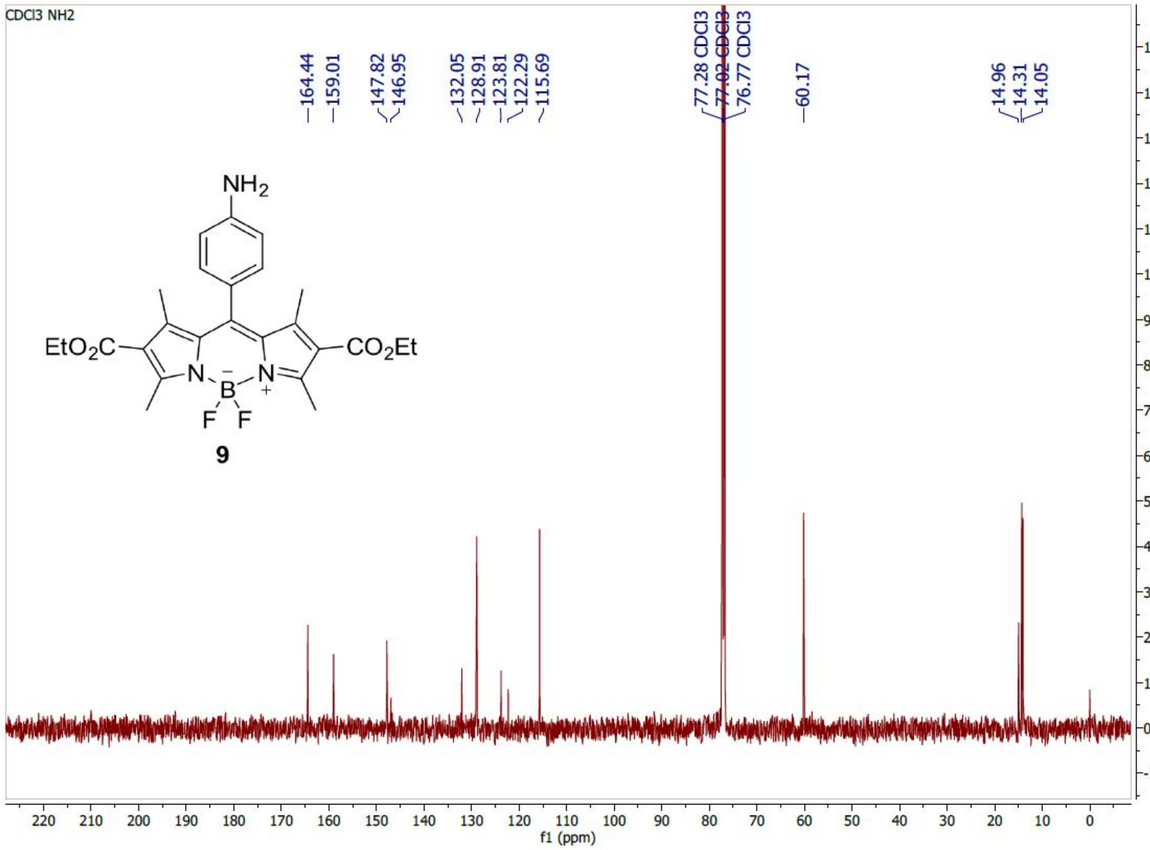

[13]C-spectrum of Diethyl 10-(4-aminophenyl)-5,5-difluoro-1,3,7,9-tetramethyl-5*H*-4l4,5l4-dipyrrolo[1,2-c:2',1'-*f*][1,3,2]diazaborinine-2,8-dicarboxylate (9)

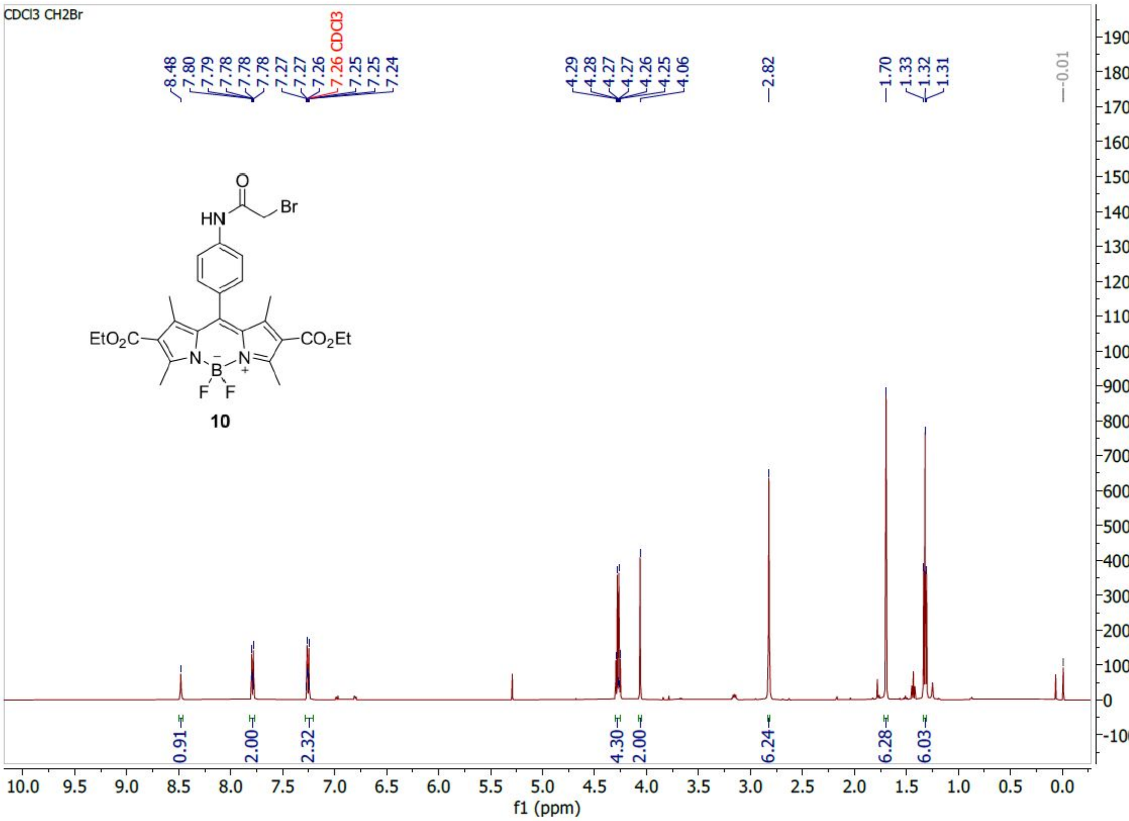

**[1]H-spectrum of Diethyl 10-(4-(2-bromoacetamido)phenyl)-5,5-difluoro-1,3,7,9-tetramethyl-5*H*-4l4,5l4-dipyrrolo[1,2-c:2',1'-*f*][1,3,2]diazaborinine-2,8-dicarboxylate (10)**

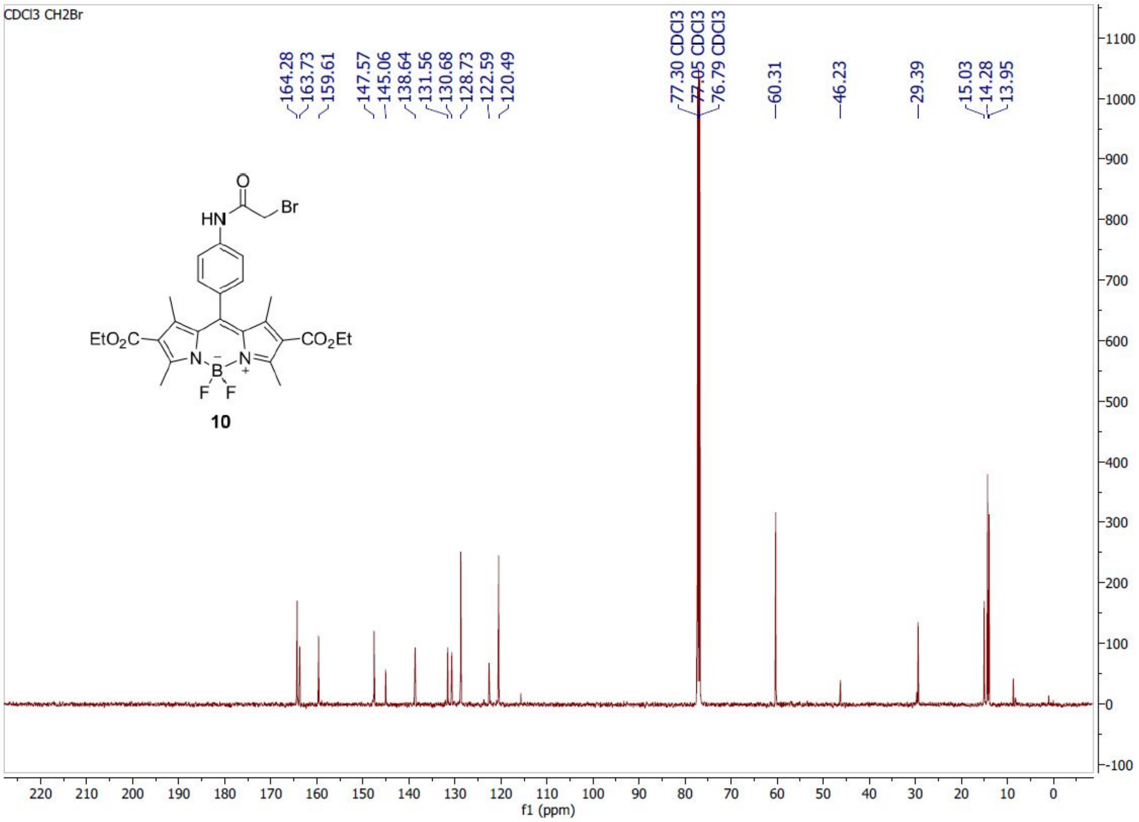

**[13]C-spectrum of Diethyl 10-(4-(2-bromoacetamido)phenyl)-5,5-difluoro-1,3,7,9-tetramethyl-5*H*-4l4,5l4-dipyrrolo[1,2-c:2',1'-*f*][1,3,2]diazaborinine-2,8-dicarboxylate (10)**

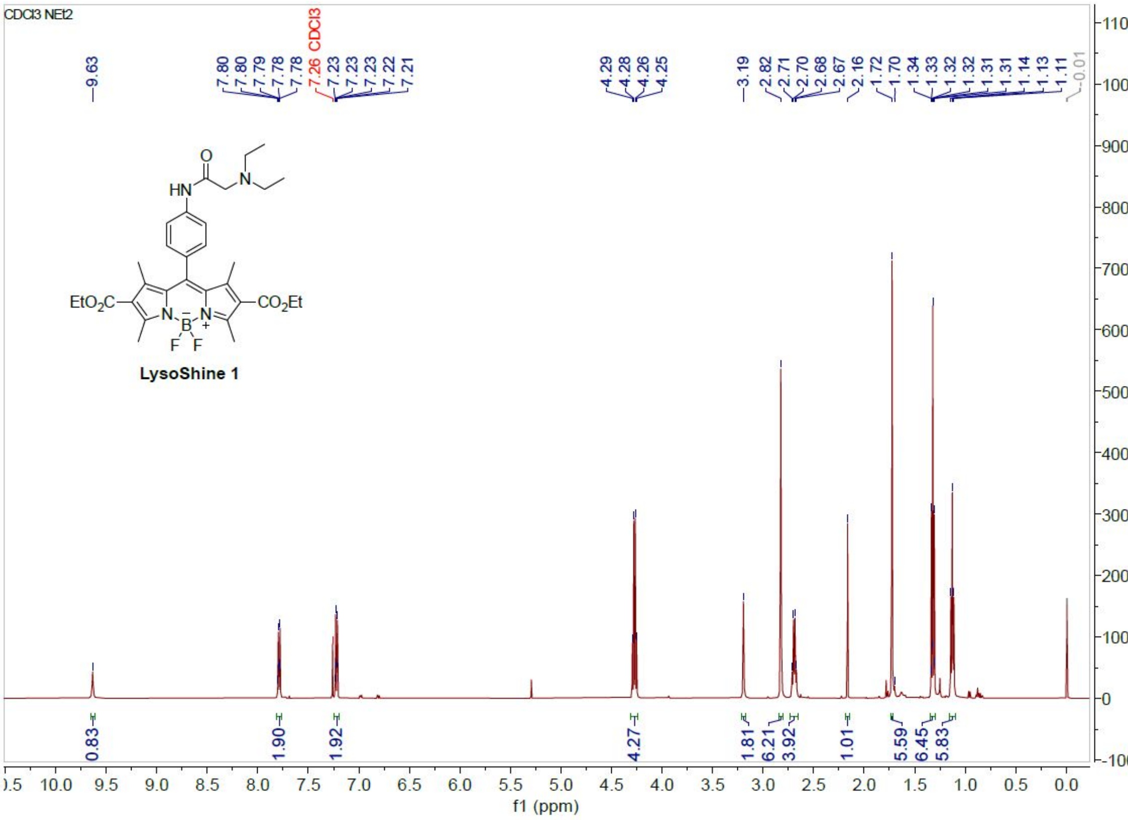

**$^{1}$H-spectrum of Diethyl 10-(4-(2-(diethylamino)acetamido)phenyl)-5,5-difluoro-1,3,7,9-tetramethyl-5*H*-4λ$^{4}$,5λ$^{4}$-dipyrrolo[1,2-c:2',1'-*f*][1,3,2]diazaborinine-2,8-dicarboxylate (LysoShine 1)**

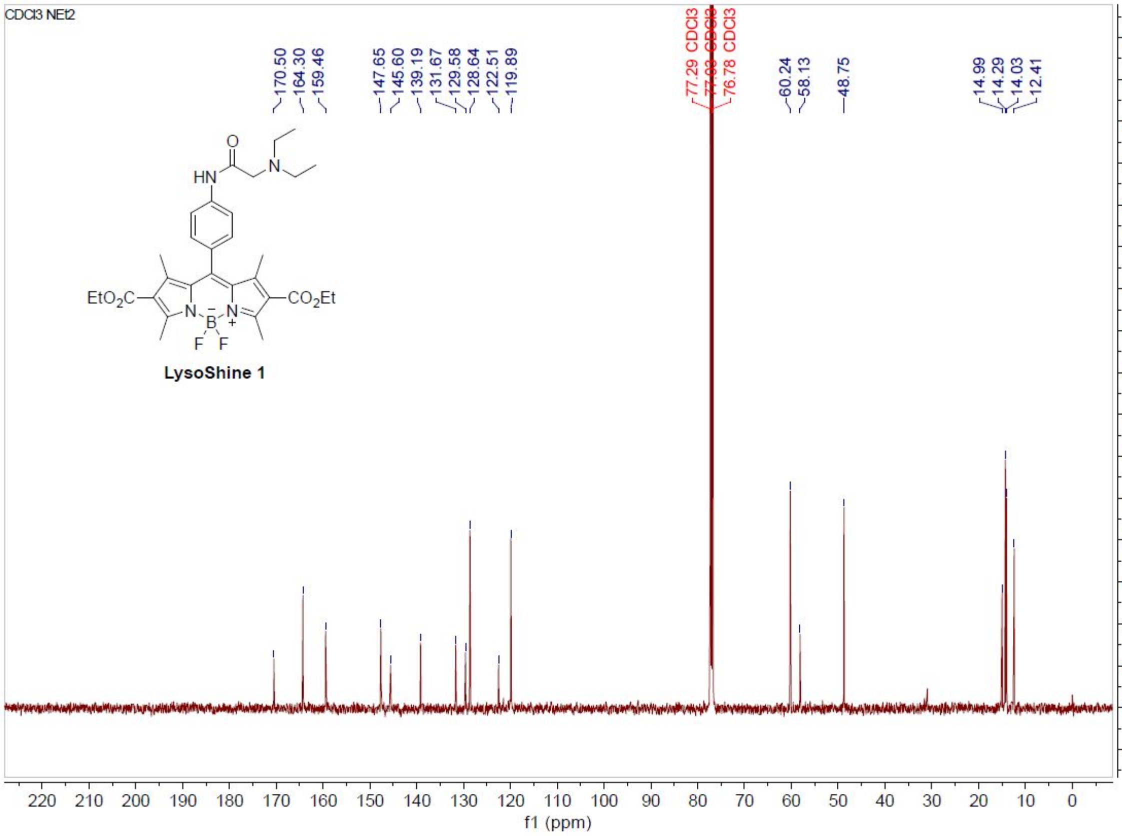

**$^{13}$C-spectrum of Diethyl 10-(4-(2-(diethylamino)acetamido)phenyl)-5,5-difluoro-1,3,7,9-tetramethyl-5*H*-4λ$^{4}$,5λ$^{4}$-dipyrrolo[1,2-c:2',1'-*f*][1,3,2]diazaborinine-2,8-dicarboxylate (LysoShine 1)**

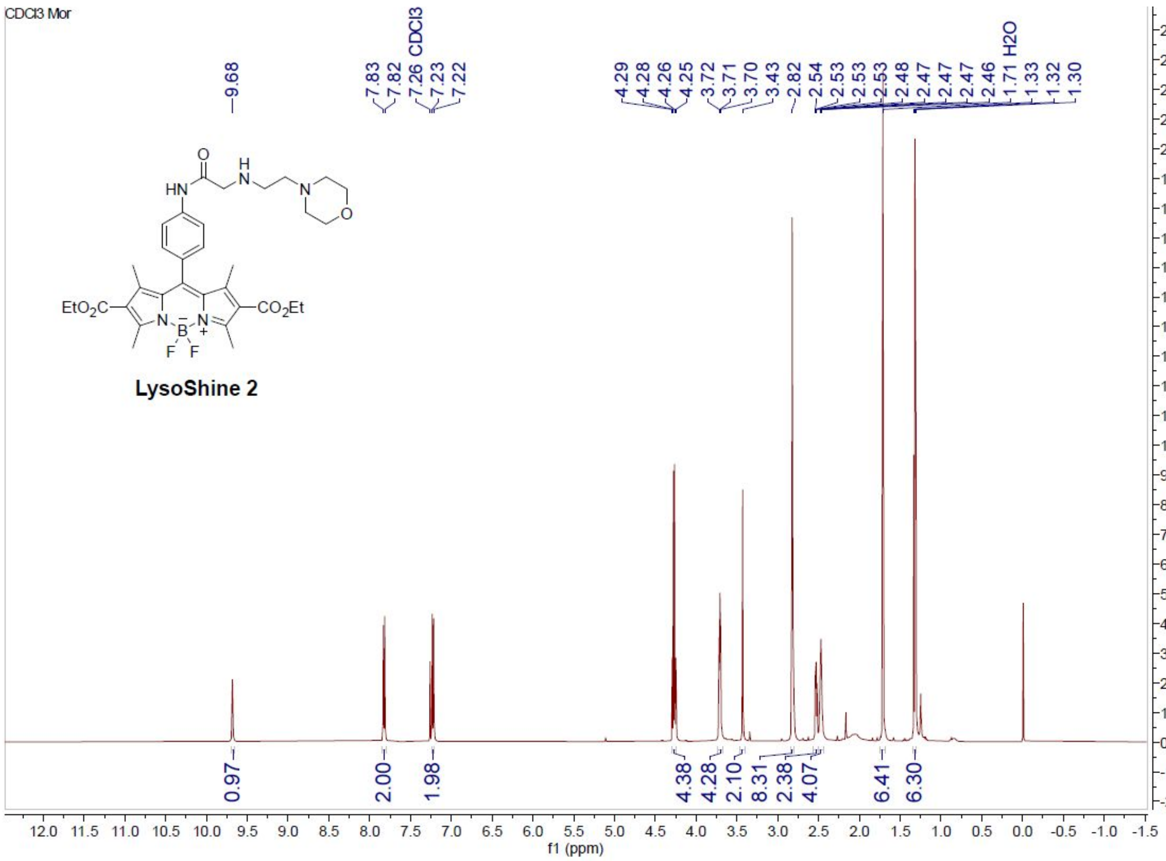

**$^{1}H$-spectrum of Diethyl 5,5-difluoro-1,3,7,9-tetramethyl-10-(4-(2-((2-morpholinoethyl)amino)acetamido)phenyl)-5*H*-4λ$^{4}$,5λ$^{4}$-dipyrrolo[1,2-c:2',1'-*f*][1,3,2]diaza-borinine-2,8-dicarboxylate (LysoShine 2)**

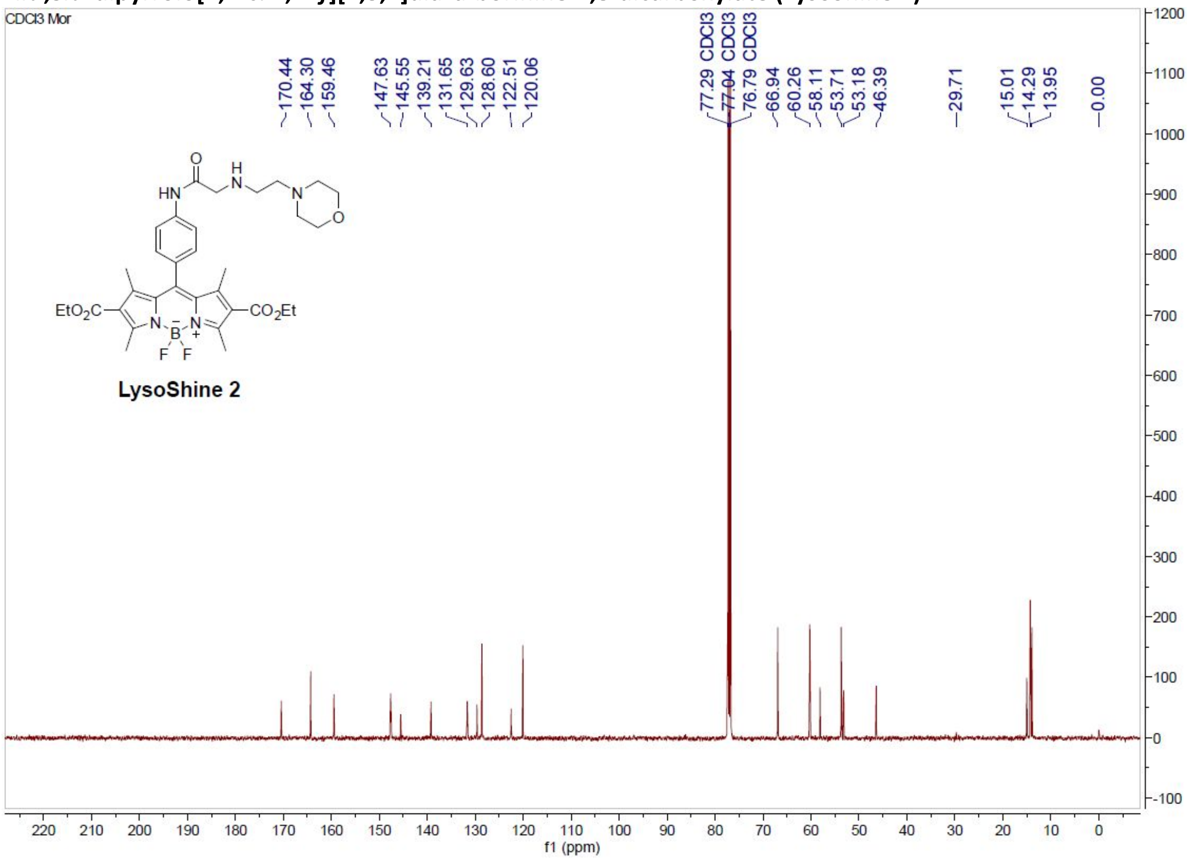

**$^{13}C$-spectrum of Diethyl 5,5-difluoro-1,3,7,9-tetramethyl-10-(4-(2-((2-morpholinoethyl)amino) acetamido)phenyl)-5*H*-4λ$^{4}$,5λ$^{4}$-dipyrrolo[1,2-c:2',1'-*f*][1,3,2]diaza-borinine-2,8-dicarboxylate (LysoShine 2)**

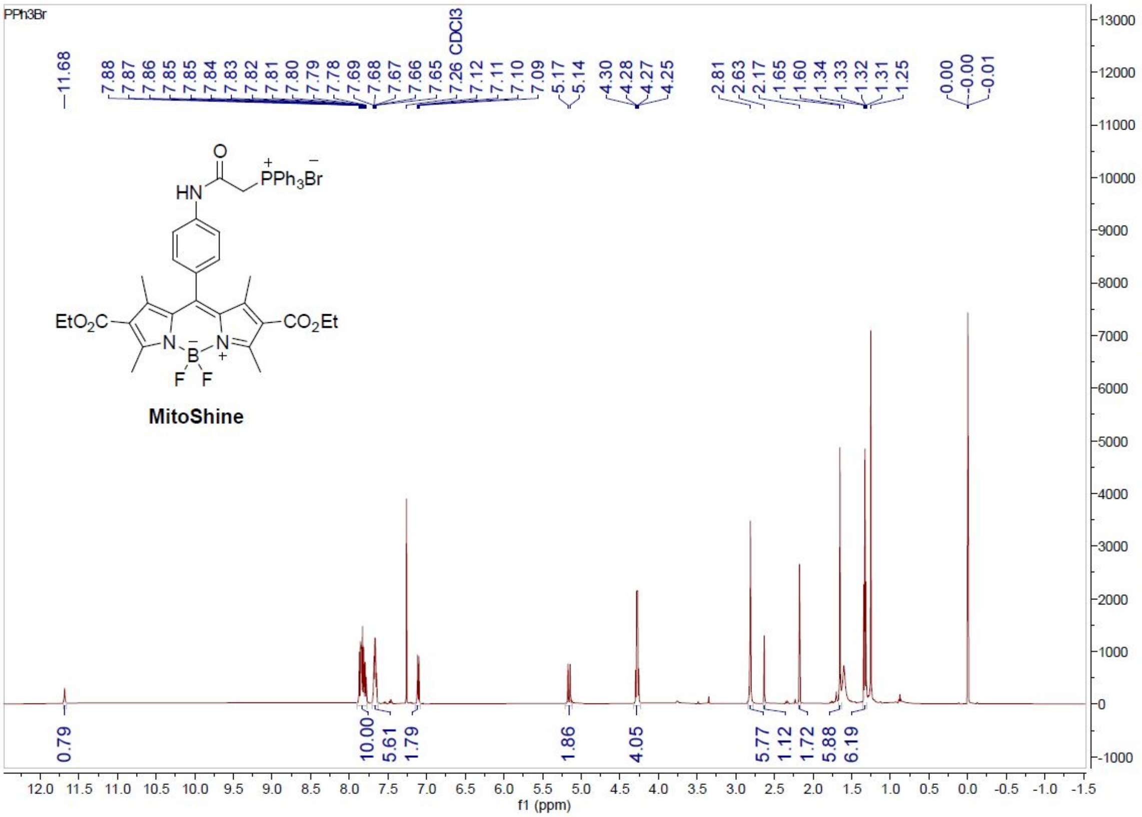

**[1]H-spectrum of (2-((4-(2,8-Bis(ethoxycarbonyl)-5,5-difluoro-1,3,7,9-tetramethyl-5*H*-4l4,5l4-dipyrrolo[1,2-c:2',1'-*f*][1,3,2]diazaborinin-10-yl)phenyl)amino)-2-oxoethyl)triphenylphosphonium bromide (MitoShine)**

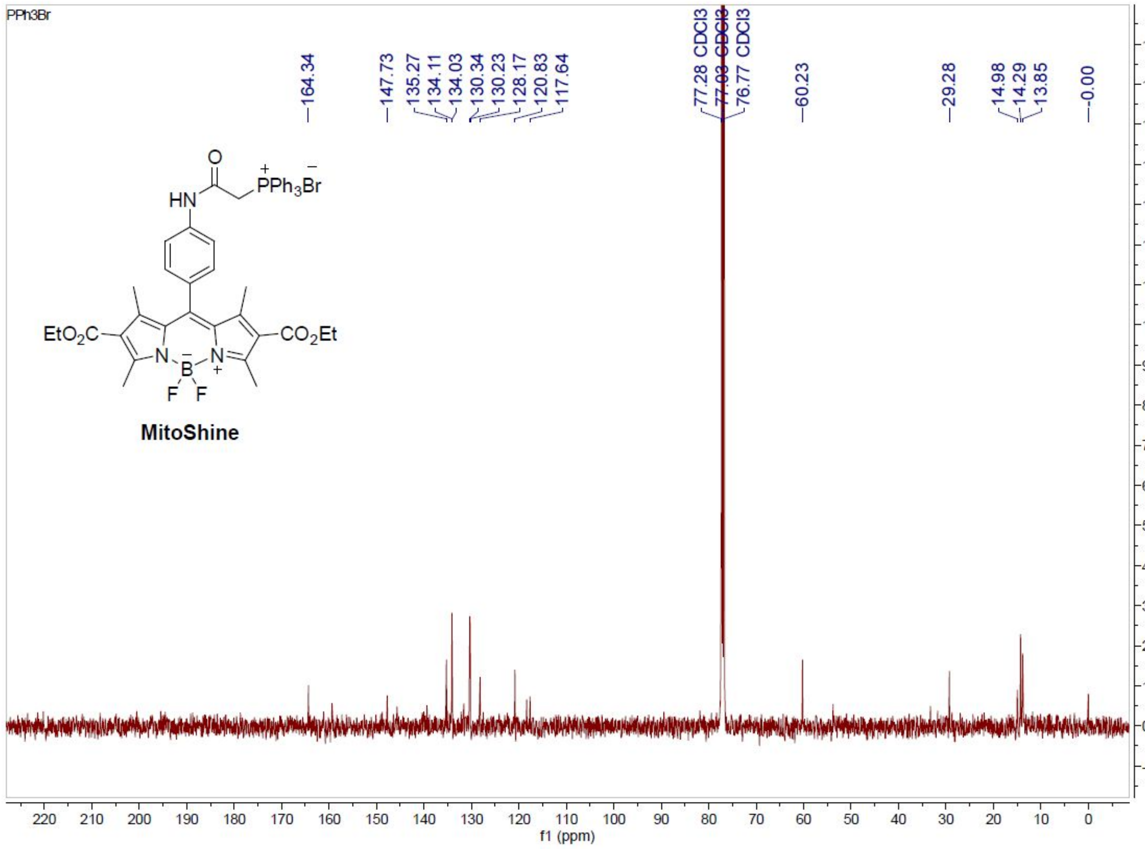

**[13]C-spectrum of (2-((4-(2,8-Bis(ethoxycarbonyl)-5,5-difluoro-1,3,7,9-tetramethyl-5*H*-4l4,5l4-dipyrrolo[1,2-c:2',1'-*f*][1,3,2]diazaborinin-10-yl)phenyl)amino)-2-oxoethyl)triphenylphosphonium bromide (MitoShine)**

#### Supporting Information References

(1) Prakash, P.; Jethava, K. P.; Korte, N.; Izquierdo, P.; Favuzzi, E.; Rose, I.; Guttenplan, K. A.; Dutta, S.; Rochet, C.; Fishell, G.; et al. Monitoring Phagocytic Uptake of Amyloid β into Glial Cell Lysosomes in Real Time. *bioRxiv* **2020**. https://doi.org/10.1101/2020.03.29.002857.

(2) Zhang, J.; Yang, M.; Li, C.; Dorh, N.; Xie, F.; Luo, F.-T.; Tiwari, A.; Liu, H. Near-Infrared Fluorescent Probes Based on Piperazine-Functionalized BODIPY Dyes for Sensitive Detection of Lysosomal PH. *J. Mater. Chem. B* **2015**, *3* (10), 2173–2184. https://doi.org/10.1039/C4TB01878H.
